# Linkage and association mapping coupled with pan-genome analyses of *Vat* homologs reveal QTLs and alleles for aphid resistance in melon

**DOI:** 10.64898/2026.04.17.719245

**Authors:** Belinchon-Moreno Javier, Coindre Eva, Monnot Severine, Berard Aurélie, Canaguier Aurélie, Le-Clainche Isabelle, Mistral Pascale, Leyre Karine, Rittener-Ruff Vincent, Hinsinger Damien D., Faivre-Rampant Patricia, Boissot Nathalie

## Abstract

Aphids threaten crop productivity through phloem feeding and the transmission of plant viruses. *Aphis gossypii*, in particular, is a widespread and damaging pest of worldwide cultivated melon. Resistance to its emerging CUC1 clone in Europe remains poorly characterized.

Here, we dissected the genetic architecture of melon resistance to CUC1 using complementary traits that capture multiple stages of the aphid-melon interaction: plant attractiveness to aphids, acceptance, aphid colonization, and multiplication. Genome-wide association studies (GWAS) in a diversity panel of 174 accessions identified a quantitative trait locus (QTL) for attractiveness on chromosome 6, while analyses in a complementary panel of 212 accessions revealed QTLs for plant acceptance by aphids on chromosomes 3, 8, and 12. Colonization and multiplication traits further highlighted resistance QTLs on chromosomes 5 and 12, the latter supported by both SNP-based GWAS and bulk-segregant analysis. Pan-NLRome k-mer- and graph-based GWAS, together with *Vat* presence/absence association analyses, provided allele-level resolution of the QTL on chromosome 5, corresponding to the *Vat* region. Leveraging allelic diversity at this locus, we functionally characterized 20 *Vat* homologs with four R65aa motifs within their leucine-rich repeat (LRR) domain and demonstrated the capacity of 4R65aa-type *Vat* alleles to confer clone-specific resistance. Resistance-conferring alleles limited virus multiplication, such as Cucumber mosaic virus (CMV), when transmitted by five *A. gossypii* clones, including CUC1.

Together, our results revealed multiple genetic determinants underlying quantitative resistance to the *A. gossypii* CUC1 clone in melon and highlighted the central role of *Vat* homologs in resistance to both *A. gossypii* and the viruses it vectors. These findings provide strategic targets for pyramiding resistance loci acting at different stages of the pest life cycle to enhance durable protection against these biotic threats.

## Introduction

Plants in agricultural environments typically encounter numerous challenges from biological agents. Among these, aphids, a diverse group of insects capable of affecting a wide range of crops ^[1]^, represent a particularly significant threat. Aphids feed on the phloem sap of vascular plants by using their piercing-sucking mouthparts, also called stylets. These stylets move through the spaces between plant cells (the apoplast) until they reach the phloem sieve elements ^[2]^. This feeding activity can severely hinder plant growth and reproduction by draining essential photoassimilates, and in some cases, by transmitting harmful plant viruses. Moreover, aphids excrete honeydew, a sugar-rich substance that, while occasionally harvested for niche products such as fir honey, typically interferes with photosynthesis and compromises fruit quality by allowing development of sooty mold ^[3]^. Developing durable plant resistance to aphids is therefore an important objective for sustainable crop protection.

Plants deploy multiple layers of defense against aphids, the first consisting in the capacity to repel them. Although plants possess a variety of mechanisms to achieve this response, their genetic basis remain poorly understood. In *Brassica juncea*, the *BjA06.GL1* and *BjB02.GL1* genes (MYB transcriptional factors) were recently found to control leaf trichome formation contributing to aphid repellence ^[4]^. Once aphids overcome this first layer of defense and feed within the phloem, plants rely on phloem-based defenses (PBD) also regulated by MYB transcriptional factors, as shown in wheat ^[5]^, chrysanthemum ^[6,7]^ and *Arabidopsis thaliana* ^[8,9]^. Additionally, members of the WRKY transcription factor family have been shown to finely regulate complex defense mechanisms and control aphid population growth in various plant species ^[10–12]^. In *A. thaliana*, the genes PHYTOALEXIN DEFICIENT4 (PAD4) and induced lipase 1 (MPL1) have been found to regulate feeding by *Myzus persicae* ^[13–16]^. All these plant genes have been identified through targeted mutant analyses and RNA expression studies; however, the extent of available natural diversity, as a resource to enhance species resistance, remains largely unexplored.

Beyond the activation of basal defense genes, Nucleotide-binding domain Leucine-rich repeat (NLR) receptor proteins constitute a distinct layer of immunity, conferring resistance to specific aphid biotypes and, in some cases, to the viruses they transmit ^[17]^. Actually, there are two principal mechanisms of virus transmission by aphids. In the non-persistent mode, virus particles are acquired during brief probing of infected plants, adhering to the aphid’s mouthparts. These particles are then quickly transmitted to new plants during subsequent feeding attempts but are also rapidly lost. The spread of non-persistent viruses can often be halted by a hypersensitive response triggered by aphid effector–mediated immunity, which is usually associated with NLR gene activity ^[18]^. On the contrary, some other viruses are persistent in aphids. In these cases, virus particles are ingested during sustained feeding on infected plants, circulate within the aphid’s body, and accumulate in the salivary glands. Transmission to new host plants then occurs specifically via the fascicular phloem, requiring full plant acceptance (successful establishment) by the aphid. Epidemics of such viruses can be mitigated when aphids show reduced acceptance of host plants, as observed in melon ^[19]^ or in cucumber ^[20]^.

Our study particularly focused on plant resistance to *A. gossypii*, a polyphagous aphid species that colonizes a wide range of plant families, including economically important crops such as malvaceous species, cucurbits, and citrus. Despite its broad host range, effective colonization of crops such as cotton or cucurbits is restricted to a limited number of clones ^[21]^. Consequently, *A. gossypii* populations in agricultural environments can comprise genetically differentiated groups ^[22]^, and the outcome of plant-aphid interactions can depend strongly on aphid genotype. This clone specificity poses a major challenge for resistance breeding: resistance effective against one aphid clone may provide limited protection against another while resistance to aphids is one of the criteria for inclusion of melon varieties in the official European and French catalogs.

Melon provides a particularly relevant system in which to investigate this problem. *A. gossypii* is an important pest of melon, causing direct damage through feeding and contributing to the spread of plant viruses. Among the few NLR genes identified to confer aphid resistance, the *Vat* gene, identified in the melon accession PI 161375 and making part of a highly diversified NLR cluster ^[23]^, is a unique model because it also confers resistance to viruses inoculated by *A. gossypii* ^[24]^. However, the long-term durability of *Vat*-based resistance is threatened by both the adaptation of aphid populations and the strong clone specificity of the resistance response. Demo-genetic analyses of *A. gossypii* populations on *Vat* and non-*Vat* melon lines during cropping seasons have identified two key bottlenecks in the maintenance of *Vat*-adapted aphid clones: low production of dispersal morphs and overwinter extinction. Both bottlenecks were observed in southwestern France, only one in southeastern France, and none in the Lesser Antilles ^[25]^, three major French melon production basins. Therefore, there is a growing interest in identifying natural genetic variation underlying aphid resistance in melon to support breeding strategies aimed at sustainable control of new aphid clones and the viruses they transmit.

While the *Vat-1* gene from the melon accession PI 161375 shows strong resistance against the NM1 *A. gossypii* clone, its effectiveness is partial or negligible against others, and quantitative resistance to *A. gossypii* has been poorly studied ^[24]^. Among the clones detected in Europe, CUC1 is of particular interest because it has been reported at high frequency in European *A. gossypii* populations ^[22]^; in our experimental work, all *A. gossypii* colonies characterized so far on *Vat*-melon in the south of France have been identified as CUC1. The heritability of its resistance, however, remains unknown.

Here, we investigated the genetic architecture underlying melon resistance to the CUC1 clone of *A. gossypii*, including the identification of quantitative trait loci (QTLs), their positions, and their additive effects. We integrated results from different projects conducted over several years to investigate novel resistance-related traits, and combined complementary genetic approaches to progressively explore the genetic basis of resistance at increasing levels of resolution. We first used bulk segregant analysis (BSA) in a bi-parental population specifically designed to avoid the confounding effect of the major *Vat*-mediated resistance. This cost-effective approach allowed us to identify genomic regions associated with resistance to colonization, an integrative trait reflecting both aphid acceptance and subsequent multiplication, inherited from a single accession. We then extended the analysis to a broader melon diversity through genome-wide association studies (GWAS) based on SNPs to dissect the genetics basis of different components of the melon–CUC1 interaction, including plant attractiveness, aphid acceptance and multiplication. Finally, we applied targeted GWAS based on k-mers and graph nodes derived from a pan-NLRome to investigate genetic associations within NLR-containing regions at higher resolution and to capture a broader representation of NLR diversity than can be obtained from a single-reference genome approach. Moving beyond resistance to the CUC1 clone, virus transmission analyses with five *A. gossypii* clones allowed us to decipher the role of multiple *Vat* homologs in clone-specific resistance. Altogether, these complementary approaches provided a genome-wide view of the genetic architecture of CUC1 resistance in melon at increased resolution, while also identifying novel *Vat* alleles conferring specific resistance to several aphid clones.

## Material and methods

### Plant materials

#### Melon panel selection for genetic association studies

We used three partially overlapping melon diversity panels (Figs. 1A and S1A, Table S1). These panels were assembled for different projects over several years, selecting accessions according to the specific objectives of each experiment and the sequencing data available at that time. Seeds were provided by the INRAE Center for Vegetable Germplasm (CRB-Leg) in Avignon ^[26]^.

**Fig. 1.**
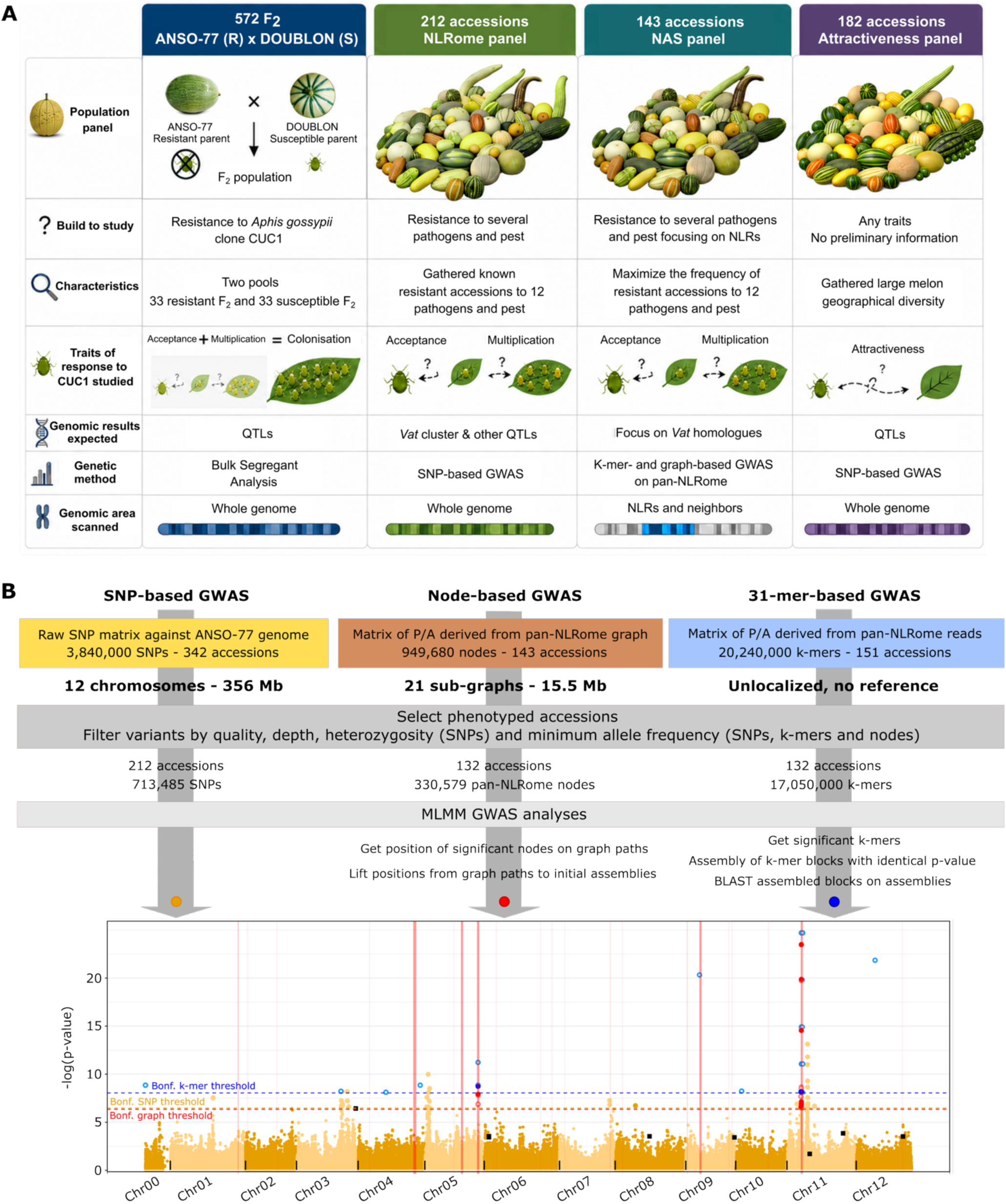
Overview of the study design and GWAS workflow. **A**, Summary of the populations and diversity panels used, including the rationale for their assembly, their main characteristics, and their utility. **B**, Summary of the GWAS workflow adapted from Belinchon-Moreno et al. ^[27]^ and applied to aphid CUC1 colonization data. SNP-based GWAS was performed at the whole-genome scale, enabling the screening of the entire chromosomes. In parallel, pan-NLRome graph node- and k-mer-based GWAS were performed across 21 targeted genomic regions containing NLRs, allowing a more detailed investigation of these regions. In the Manhattan plot, orange dots represent SNPs tested in the first MLMM step, whereas black squares represent SNPs highlighted in subsequent MLMM steps. Black vertical lines delimit chromosomes, and red vertical bars locate NLR-containing genomic regions, with their widths scaled according to region size. Significant graph nodes (red dots) and k-mers (blue dots) are overlaid on the corresponding genomic regions. K-mers and graph corresponding to sequences absent from the reference genome are shown as lighter, unfilled dots. Absent k-mers were forced to align to the region with the highest sequence similarity, whereas absent graph nodes were positioned on the closest graph node containing a reference genome sequence.

The first panel (PA_colonization_, 212 accessions) was used to evaluate aphid colonization, a trait including both the initial acceptance of the plant by aphids and the subsequent ability of aphids to multiply on the plant. This panel was originally assembled to investigate melon resistance to multiple pests and pathogens, not only aphids, and has already been characterized and used in previous studies ^[27]^. The second panel (PA_attractiveness_, 182 accessions, included 87 shared with PA_colonization_) was used to evaluate plant attractiveness to aphids. Among the accessions included, 181 belonged to the *C. melo* species, and one was part of the *C. picrocarpus* species. We assembled this panel with the objective of investigating whether variation in melon attractiveness to *A. gossypii* could be genetically controlled independently of the major immune-related loci already known to contribute to aphid resistance. Therefore, we tried to maximize the geographical and genetic diversity of melon accessions for which sequencing data was available. The third panel (PA_NAS_, 143 accessions of which 132 were common to the colonization panel and 35 to the attractiveness panel) was completely sequenced using Nanopore adaptive sampling (NAS) to selectively enrich genomic regions containing NLR genes using long reads, as described in Belinchon-Moreno et al. ^[27]^. We used this PA_NAS_ diversity panel to focus on resistance encoded on NLR regions, and therefore we tried to maximize the representation of resistant accessions.

Together, these panels captured a worldwide diversity and encompassed accessions from 17 different botanical groups (Table S1). They contained ancestral recombination allowing to develop genetic associations studies based on linkage disequilibrium.

#### Generation of bi-parental populations

##### ANSO-77 x VEDRANTAIS

We produced 186 F_2_ plantlets from a cross between ANSO-77 (Spanish accession, botanical group *inodorus*, resistant to *A. gossypii* clone CUC1 colonization) and VEDRANTAIS (French line, botanical group *cantalupensis*, susceptible to *A. gossypii* clone CUC1 colonization), to assess whether the *Vat* gene cluster ^[28]^ of ANSO-77 carries the resistance to colonization by the CUC1 clone of *A. gossypii*. We extracted DNA from the 186 F_2_ plantlets as described in the following section and amplified by PCR four dominant markers distinguishing the parental *Vat* haplotypes: Z5259F (TTGTGGAAGATTGAGTAGTTTAACTT) and Z5238R (CAAGAAGTAATGGTTAAATTGCGTAGT), specific to ANSO-77, and Z1431F (ATGCAAAGAGTTTGAAGATG) and Z5239R (CATTCAGAAATGGATACATCAG), specific to VEDRANTAIS. PCR reactions were carried out in two 96-well plates including both parental lines and F_1_ hybrids as controls. We selected 38 F_2_ presenting the same amplification pattern as VEDRANTAIS, and 49 presenting that of ANSO-77. These 87 plantlets were phenotyped for resistance to colonization by *A. gossypii* clone CUC1 as described below.

##### ANSO-77 x DOUBLON

We derived 576 F_2_ plants from a cross between ANSO-77 and DOUBLON (French line, botanical group *cantalupensis,* highly susceptible to *A. gossypii* clone CUC1 colonization) and we performed a Bulk Segregant Analysis (BSA) with the objective of identifying genomic regions associated with resistance to CUC1 beyond *Vat*. We self-crossed each F_2_ to obtain F_3_ families, which were first phenotyped for colonization by *A. gossypii* clone CUC1 using mass inoculation bioassays (see phenotyping procedures) to eliminate families showing intermediate levels of aphid colonization. We selected 202 F_3_ families presenting extreme scores, and we evaluated their response to aphid colonization using an inoculation by individual plant (see phenotyping procedures). We inferred the resistance or susceptibility of each F₂ plant to aphid colonization from the phenotype of its F₃ progeny. For each selected F_3_ family, we calculated a normalized colonization score relative to resistant and susceptible controls, using equation (1),

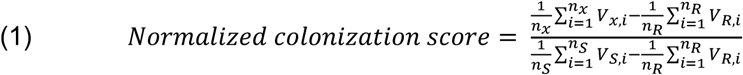

where 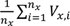 is the average colonization score of F_3_ plants belonging to family 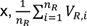 is the average colonization score of resistant (R) controls present in the tests were F_3_ plants from the family x where scored, and 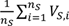 is the average colonization score of susceptible (S) controls present in the tests where F_3_ plants from the family x where scored.

Based on the normalized score of the 202 F_3_ families, we selected two contrasting bulks of 38 families: a resistant bulk with the lowest colonization levels, and a susceptible bulk with the highest colonization levels.

### DNA extraction and sequencing

#### Short-read sequencing

We obtained short-read whole-genome sequences for 292 accessions from the PA_colonization_ and PA_attractiveness_ panels using Illumina and/or MGI technologies, achieving genome coverages ranging from 13× to 78× (Table S1). Complementarily, we retrieved short reads for 15 accessions from NCBI to complete the panels (Table S1). Furthermore, we produced whole-genome Illumina sequencing data for the parental lines and for the R and S F₂ bulks from the ANSO-77 × DOUBLON population (Table S1).

We sampled 150 to 200 mg of young plant leaves and froze them in liquid nitrogen for DNA extraction. We extracted DNA with the Qiagen DNAeasy Plant miniKit (Qiagen, Valencia, CA, USA) and evaluated DNA quantity and quality as described in Belinchon-Moreno et al. ^[27,29]^. For bulks sequencing, DNA from individual F₂ plants classified as R and S was pooled in equimolar concentrations and sequenced separately. We prepared sequencing libraries as presented in Belinchon-Moreno et al. ^[29]^, using the Kapa Hyper Prep Kit PCR-Free (KapaBiosystems, Wilmington, MA, USA) for Illumina libraries, and the MGIEasy PCR-Free DNA Library Prep Kit (MGI, Shenzhen, China) for MGI libraries. We sequenced Illumina libraries on an Illumina NovaSeq6000 instrument (Illumina, San Diego, CA, USA) setting the 150 bp paired-end reads configuration. We loaded MGI libraries onto an MGI DNBSEQ-G400RS device (MGI, Shenzhen, China) with G400HM flowcells and HotMPS High-throughput Sequencing chemistry to generate paired-end 150 bp reads. Raw Illumina reads were processed following Alberti et al. ^[30]^.

#### Long-read sequencing and NLRome assemblies

We retrieved NAS sequencing reads for the 143 accessions composing the PA_NAS_ panel from Belinchon-Moreno et al. ^[27]^. These reads were generated to enrich genomic regions containing NLR genes, typically characterized by their high number of copy number variations (CNVs), presence-absence (P/A) polymorphisms and repetitive elements. We also recovered the NLRome assemblies generated from NAS-sequencing reads from Belinchon-Moreno et al. ^[27]^. Briefly, these NLRome assemblies incuded 15 *de novo*–assembled NLR regions enriched by NAS, and six isolated NLR genes or domains obtained through guided assembly.

### Phenotyping procedures and data analysis

#### Aphid rearing and maintenance

*A. gossypii* clones collected from cucurbits were reared on melon VEDRANTAIS plants at 24/18 °C (day/night) under a 16h/8h light/dark photoperiod. The *A. gossypii* clones CUC1, NM1, C9 (collected in France), GWD2 and C6 (collected on Lesser Antilles) were previously characterized using 15 microsatellite markers ^[22]^. We used apterous adult aphids of five-to-seven days old for carrying out the biological tests.

#### Plant attractiveness to aphids

We tested the 182 accessions composing the PA_attractiveness_ panel for their attractiveness to *A. gossypii* clone CUC1 in a multiple-choice bioassay as described in Monnot et al. ^[20]^. In this bioassay, aphids were allowed to freely select among genotypes ^[31]^. For each tested genotype, we collected nine leaf disks of 2 cm diameter from four 21-day-old plantlets and placed them with their upper leaf side facing up on agar medium (4 g/L agar, 10 g/L mannitol) in a closed box, forming a row (Fig. S2). For a given accession, leaf disks were collected from as many plants as possible to capture intra-genotype diversity. Each agar-filled box contained two rows of two randomly chosen genotypes, flanked by a row of disks from VEDRANTAIS (attractive control) and a row of disks from VIRGOS (repulsive control) (Fig. S2). We deposited one apterous aphid five-to-seven days old on each leaf disk from the two tested genotypes (18 aphids per box). We stored boxes for 24 hours in a climate chamber at 24/18 °C (day/night) with a photoperiod of 16/8h light/dark. We counted the number of aphids feeding on each tested genotype on the following day. We ran six independent assays (330 boxes), allowing each genotype to be observed in at least four different boxes. We hypothesized that the higher the number of aphids feeding on a genotype, the more attractive this genotype was for aphids. To quantify relative attractiveness, we modeled aphids counts using a multinomial likelihood framework, assuming: i/ Aphid behavior is statistically identical within a box; and ii/ Aphids acted independently. To simplify the model, we assumed that attractiveness depends only on the genotype of each disk.

Let *a*_g*i*_ denote the attractiveness of the genotype *g_i_*, and *r*_g*i*_ the number of disks of genotype *g_i_* in a given box *b*. Let *G_b_* be the set of genotypes present in the box *b*, including *g_i_* itself, the other tested genotype in the same box, as well as resistant or susceptible controls. Then, the probability that a randomly chosen aphid occupies a disk of genotype *g_i_* is given by equation (2).

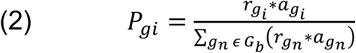

Consequently, the number of aphids *n*_g_*_i_* observed on genotype *g_i_* in a box containing *N_b_* aphids follow a multinomial distribution with parameters (*N_b_*, *P*_g1,_ …, *P*_g*G*_), where *G* is the number of genotypes in the box. The likelihood of observing the data in a given box *b* is the product of probabilities across all genotypes present in that box (*G_b_*), including resistant and susceptible controls, as described by equation (3):

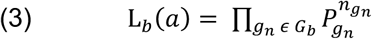

and the overall likelihood across all boxes, assumed independent, is then described by equation (4):

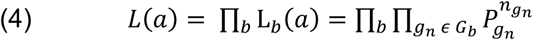

To remove indeterminacy due to the relative scale of attractiveness, we constrained the sum of genotype attractiveness values across all the genotypes included in the panel (*G_P_*) to 1: ∑_g_*_n_εG_P_* (*a*_g_*_n_*) = 1. Maximum likelihood estimation of *a*_g_*_i_* was performed using numerical optimization of the negative log-likelihood, reparameterizing *a*_g_*_i_* as in equation (5), to ensure positivity and normalization:

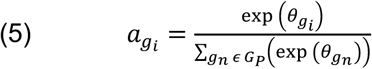

To quantify uncertainty in the estimates, we simulated 1,000 datasets using the estimated multinomial model, re-estimated *a*_g*i*_ for each simulation, and derived 95% confidence intervals from the 2.5th and 97.5th percentiles. We normalized these data using the bestNormalize R package ^[32]^, prior to GWAS analyses. The selected normalization was log_10_(x + 0.0009999964).

Finally, we calculated a heritability-like measure of attractiveness, defined as the proportion of variance in estimated attractiveness explained by genotype, corrected for the estimation of variance, as described in equation (6),

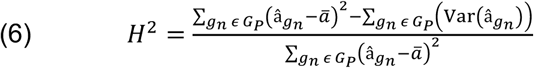

where â_g_*_n_* is the estimated attractiveness of genotype *g_n_* and a̅ is the mean across all genotypes. This whole approach allowed us to quantify relative attractiveness of each genotype to aphids, assess statistical confidence, and estimate the genetic contribution to aphid preference.

#### Plant resistance to aphid colonization, including acceptance and multiplication

We assessed acceptance of the plant by *A. gossypii* and their ability to restrict aphid colonization, as described by Thomas et al. ^[33]^. Briefly, we deposited 10 apterous adults on each plantlet and grew them in a climate chamber at 24/18 °C (day/night) with a photoperiod of 16/8h light/dark. Three days after inoculation, we recorded the number of aphids remaining on the plantlet as the acceptance phenotype. Aphids that did not accept the plant left the plant. Seven days after inoculation, we recorded the number of adult aphids still present on each plantlet and estimated nymph abundance using a 0-6 scale (0: no nymphs; 1: 1-20 nymphs; 2: 20-50 nymphs; 3: more than ∼50 nymphs observed on the infested leaf and elsewhere on the plantlet; the two scores were summed). We calculated a colonization index as [nymph density + ln(number of adults + 0.001)].

For the 576 F_3_ families derived from the ANSO-77 x DOUBLON cross, we performed a first-stage phenotyping using mass inoculation bioassays to discard intermediate phenotypes. We used plastic trays, each containing 60 plants (10 × 6 grid). We included two tested genotypes per tray (20 seeds per genotype, sown on outer rows), with the two central rows reserved for resistant (VIRGOS) and susceptible (VEDRANTAIS) controls. At least 18 plantlets developed for 475 families and were scored ten days after inoculation, following a notation as very resistant, resistant, intermediate-resistant, susceptible, and very susceptible, based on the global response of the family. Seventy-five F_3_ families were classified as very resistant or resistant, while 77 as very susceptible. For the 323 F_3_ families classified as intermediate-resistant or susceptible, we performed a second phenotyping step with an identical configuration, seeding in this case 30 seeds per F_3_ family (15 seeds per row). Of the 281 F_3_ families with ≥18 individuals, 29 were classified as resistant, and 99 as susceptible or very susceptible. Altogether, 280 F_3_ families showed extreme phenotypes (104 resistant and 176 susceptible). From these, 202 families with enough seeds available were retained for individual-plant phenotyping.

For the 87 selected F_2_ progeny (ANSO-77 x VEDRANTAIS), the 202 F_3_ selected families (ANSO-77 x DOUBLON), and all accessions of the PA_colonization_ panel, we performed a phenotyping of aphid colonization using no choice bioassays on individual plants. For the ANSO-77 x DOUBLON derived families, we implemented 50 incomplete blocks (≥15 plants for most families), always including R (VIRGOS) and S (VEDRANTAIS) controls. To phenotype the PA_colonization_ panel, we implemented 32 incomplete blocks (≥10 plants per accession).

For GWAS, we excluded observations with > 11 aphids on single plantlets (“acceptance” phenotype), and we removed genotypes with < 7 observations. We also discarded within-accession outliers (> 3 SD). We condensed up acceptance and multiplication observations on the PA_colonization_ panel to single Best-Linear-Unbiased-Predictors (BLUPs) per genotype using the mixed linear models (MLMs) presented in equations (7) and (8) for acceptance and multiplication, respectively,

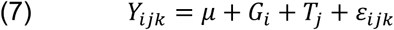

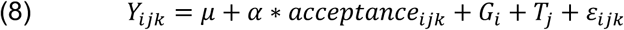

where *Y_i_*_j*k*_ denotes raw phenotypic observations; *μ* is the fixed intercept; *G_i_* is the random effect of genotype *i*, with distribution *G_i_*∼*N*(0, *Iσ*^2^); *T*_j_ is the random effect of test j, assumed to be distributed as *T_J_*∼*N*(0, *Iσ*^2^); *α* is the fixed effect of acceptance; *acceptance_i_*_j*k*_ is the measured acceptance score associated with observation *Y_i_*_j*k*_; and *ε_i_*_j*k*_ represents the residual error effect, expected to follow de distribution *ε_i_*_j*k*_∼*N*(0, *Iσ*^2^). We applied the lmer() function of the R lme4 package ^[34]^ to fit MLMs. Residuals normality was examined using Q–Q plots.

Based on Perchepied et al. ^[35]^, we estimated broad-sense heritabilities (equation 9) using the variance components from mixed models analogous to those in equations (7) and (8), but treating all factors as random.

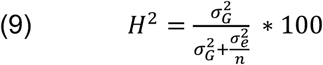

where *σ^2^_G_* is the variance associated with the genotype factor; *σ^2^_e_* is the sum of the variants of all other effects included in the model; and n is the average number of observations per genotype across the entire design (∼10.6), excluding controls.

#### Plant resistance to virus when inoculated by aphids

We ran bioassays to assess plant resistance to virus when inoculated by *A. gossypii*, as described in Thomas et al. ^[33]^. Aphids from mass rearing were allowed to feed during ten minutes on leaves of the melon cultivar VEDRANTAIS infected with CMV isolate I17F to acquire the virus. Afterwards, we deposited batches of ten aphids of a given clone (CUC1, NM1, C9, GWD2 or C6) on individual plantlets for inoculation. After 15 minutes, we removed the aphids from the plantlets, we sprayed plants with an aphicide, and we subsequently transferred them to an insect-proof greenhouse. We evaluated the occurrence of infected plants 20 days after inoculation by visual assessment of symptoms.

Phenotyping was performed in 10 incomplete blocks for C6, 33 for C9, 13 for CUC1, seven for GWD2, and 46 for NM1. We included VEDRANTAIS and VIRGOS plants in each incomplete block as susceptible and resistant controls, respectively. We tested five to 169 plantlets (average of 29) per accession and aphid clone.

### Pre-mapping approaches

#### SNPs and short INDELs matrix for Bulk Segregant Analysis

We applied the whole BSA with Next-Generation Sequencing (BSA-NGS) using the Snakemake pipeline easy_qtlseq (https://forge.inrae.fr/gafl/pipelines_snakemake/easy_qtlseq), which integrates short-reads mapping and variant matrix generation. Briefly, we processed adapters-removed and trimmed reads from the parental lines and for the resistant and susceptible F₂ bulks of the ANSO-77 × DOUBLON population with fastp v. 0.20.0 ^[36]^, merging overlapping read pairs and filtering out reads < 50 bp. Then, we mapped processed reads to the genome assemblies of ANSO-77 and DOUBLON ^[37]^ using bwa mem v. 0.7.17 ^[38]^. We removed PCR duplicates using sambamba markdup v. 0.7.1 ^[39]^ including the option --remove-duplicates. Subsequently, we applied GATK4 v. 4.1.4.1 ^[40]^ for SNP and INDEL calling on both parental genomes. We hard-filtered the raw variant callset using GATK Best Practices (https://gatk.broadinstitute.org/hc/en-us/articles/360035890471-Hard-filtering-germline-short-variants) with the modifications “FS_filter > 200” and “ReadPosRankSum_filter < −20” for INDELs filtering. We removed INDELs longer than 10 bp, and we retained biallelic variants without missing genotypes. We further retained variants with a minimum sequencing depth (DP) of five from the parental lines, and a DP between 70 and 1,400 from both F_2_ bulks. Afterwards, we kept variants that presented a polymorphism between both parental lines, and restricted the dataset to R and S bulks. This resulted in a matrix with 665,603 variants when mapping to ANSO-77 (587,287 SNPs and 78,316 INDELs ≤ 10 bp). When mapping to DOUBLON, we identified 677,435 variants (597,698 SNPs and 79,737 INDELs).

#### SNP matrices for GWAS

We used a SNP matrix from short-reads data of 342 accessions available at INRAE-GAFL, including those used in this study. This matrix was constructed using the genome assembly of ANSO-77 ^[37]^ as reference for read mapping and variant calling (Fig. 1). Only biallelic SNPs were retained, and eight accessions (ME00605, ME00470, ME00721, ME01336, ME02964, ME00232, ME02619, and ME02584) were filtered out as they presented low genotyping quality and sequencing depth. The whole process was implemented using the wgs_gatk Snakemake pipeline available at https://forgemia.inra.fr/gafl/pipelines_snakemake/wgs_gatk.

Using bcftools v.1.19, we extracted two subsets from the raw SNP matrix: one comprising 226 accessions (hereafter MA_set1_), including the 212 accessions in PA_colonization_; and the second comprising 174 accessions (hereafter MA_set2_), corresponding to all accessions in PA_attractiveness_, except for the eight excluded by their low quality. Within each subset, we set genotype calls with a genotype quality (GQ) < 15 or a DP < 5 as missing. Then, we removed variants that presented missing data, GQ < 20 or DP < 7 in more than 3% (MA_set1_) or 4% (MA_set2_) of the samples. We also discarded variants with a heterozygosity rate exceeding 50%.

From the filtered MA_set1_, we removed variants with a minor allele frequency (MAF) under 5% (∼10 accessions) and selected 212 accessions phenotyped for colonization, resulting in a SNP matrix called MA_colonization_ that included 713,485 SNPs, used for GWAS. One additional subset was defined from the filtered MA_set1_ by selecting 98 accessions phenotyped for colonization, excluding those carrying *Vat* homologs likely associated with *A. gossypii* CUC1 resistance. After filtering out variants with MAF < 6% (∼6 accessions), this selected matrix contained 599,886 SNPs.

For the filtered MA_set2_, we applied a MAF of 0.06 (∼10 accessions), producing a SNP matrix (MA_attractiveness_) that contained 174 accessions and 533,109 SNPs.

#### Pan-NLRome k-mer and node matrices from NAS reads

We used P/A matrices of k-mers and pan-NLRome graph nodes retrieved from Belinchon-Moreno et al. ^[27]^. These P/A matrices were constructed as follows:

Shortly, 31-mers were extracted from NLR regions across 151 NAS-sequenced accessions ^[27]^ using a custom Snakemake pipeline (https://forge.inrae.fr/gafl/users/javier_belinchon/prekmergwas) based on the approach developed by Voichek and Weigel ^[41]^. NAS reads from 15 NLR-rich targeted regions, along with a combination of raw NAS and short reads from six additional regions containing isolated NLRs, were provided to the pipeline. Then, kmers GWAS v. 0.3 ^[41]^ was used to consolidate k-mers count per accession, convert the list of k-mers into a P/A matrix, and compute a kinship matrix based on k-mer counts.

The retrieved k-mer P/A matrix comprised 151 accessions and ∼20.24 million k-mers. From this matrix, we retained 132 accessions whose NLRomes were fully assembled ^[27]^ and for which phenotypic data on *A. gossypii* acceptance and multiplication were available. We further applied a MAF cutoff of 0.06, resulting in ∼17.05 million k-mers used for GWAS.

The retrieved P/A matrix of pan-NLRome graph nodes was built from a pan-NLRome graph comprising 21 sub-graphs across 143 accessions ^[27]^. Each sub-graph represented one NLR-rich genomic region, and the whole graph presented a total size of 15.5 Mb, with 949,690 nodes and 1,314,840 edges. This graph was constructed using PGGB ^[42]^, and the whole process is detailed in Belinchon-Moreno et al. ^[27]^. A P/A matrix containing 143 accessions and 949.68K variant nodes was generated from this graph ^[27]^. We retrieved this matrix, and we retained 132 accessions for which phenotypic data on *A. gossypii* acceptance and multiplication were available. We then applied a MAF cutoff of 0.06, resulting in 330,579 variant nodes used for GWAS.

#### Presence/absence matrix of Vat alleles from NAS assemblies

We constructed a P/A matrix of non-redundant *Vat* alleles across accessions starting from pan-NLRome assemblies and NLR annotations retrieved from Belinchon-Moreno et al. ^[27]^. We selected *Vat* alleles across the 143 assembled NLRomes, and we clustered them at the protein level, considering as non-redundant those alleles with different protein sequences (at least one amino-acid different). The generated P/A matrix contained 143 accessions and 167 non-redundant *Vat* alleles. From this matrix, we retained 132 with available phenotypic data on *A. gossypii* acceptance and multiplication, and we applied a MAF cutoff of 0.03 (allele present in at least four accessions), yielding a matrix with 32 *Vat* alleles for GWAS.

#### Population structure and kinship assessment

We assessed population structure for the PA_attractiveness_ panel using the MA_attractiveness_ SNP matrix (174 accessions) and for the PA_colonization_ panel using the filtered MA_set1_ SNP-matrix (226 accessions, including the 212 in PA_colonization_) with a MAF cutoff of 0.05. We reduced marker redundancy by selecting 8,000 independent SNPs through linkage disequilibrium (LD) pruning in Plink v. 1.90 ^[43]^, using a sliding window of 100 kb, step size of one SNP, and r² < 0.2. We then inferred population structure with STRUCTURE v. 2.3.4 ^[44]^, testing 1-10 groups, with 10 independent runs per K value. Each run included 250,000 burn-in iterations followed by 500,000 iterations of the Markov chain Monte Carlo (MCMC). We determined optimal grouping using STRUCTURE Harvester ^[45]^, based on the Evanno method ^[46]^. We used a Snakemake workflow (https://forge.inrae.fr/gafl/users/javier_belinchon/structure) to run the whole STRUCTURE pipeline.

We calculated additive kinship on SNP and graph nodes matrices used for GWAS by applying the A.mat() function from the R package rrBLUP ^[47]^. We inferred kinship on the k-mer content across the 151 NAS-sequenced accessions ^[27]^ using kmersGWAS v. 0.3, which computed a kinship matrix directly from k-mer counts.

### Genetic mapping procedures

#### Bulk Segregant Analysis with Next-Generation Sequencing

We conducted two independent BSA-NGS analyses on the F₂ population derived from the ANSO-77 × DOUBLON cross, using the genome assemblies of each parental line as alternative references. We conducted the analyses using the Snakemake pipeline easy_qtlseq (https://forge.inrae.fr/gafl/pipelines_snakemake/easy_qtlseq), based on the QTL-seqr R package ^[48]^. Our BSA-NGS analysis employed two complementary statistical approaches, QTL-seq and G’. Both were implemented using the functions runQTLseqAnalysis() and runGprimeAnalysis() from the QTL-seqr package, following the principles described in Takagi et al. ^[49]^ and Magwene et al. ^[50]^. The function runQTLseqAnalysis() calculated the ΔSNP-index for each variant position as the difference between SNP-index (frequency of the alternative allele) of resistant and susceptible bulks ^[49]^. Tricube-smoothed ΔSNP-index across the entire genome were calculated with a sliding window of 1 Mb, smoothing out noise while accounting for LD between variants, as variants that are close to the focal variant have a high weighting value, compared to those closer to the edge of the window ^[48]^. Higher absolute ΔSNP-index values reflect stronger allele frequency differences between bulks, suggesting tight linkage to causal QTLs. We set confidence intervals of 95, 99 and 99%, which were calculated by the runQTLseqAnalysis() function by performing 10,000 bootstrapped simulations (replications), under the null hypothesis, considering a F_2_ population type (popStruc) and bulk sizes of 38 individuals (bulkSize).

For the G’ method, the function runGprimeAnalysis() first compared observed allele counts in high and low bulks to expected counts under the null hypothesis of no QTL, producing a G statistic ^[50]^. These values are then smoothed using a tricube-weighted kernel across a 1 Mb sliding window (windowSize), producing the G′ statistic and a smoothed ΔSNP-index. To assess statistical significance, the null distribution of G′ was estimated by excluding putative QTL regions and recalculating the mean and variance of the remaining SNPs. P-values were derived from the approximated G’ distribution and adjusted using the Benjamini–Hochberg false discovery rate (FDR) method. Parameters ‘outlierFilter = “deltaSNP10”’ and ‘filterThreshold = 0.1’ removed from the null distribution regions where the absolute ΔSNP-index exceeds 0.1. We pinpointed QTLs from the G’ statistics based on an FDR of 0.01.

The Snakemake pipeline also ran the function getQTLTable() to summarize genomic regions exceeding the 95% confidence interval in QTL-seq analysis, and the FDR threshold of 0.01 in the G’ method.

#### SNP, k-mer and graph node GWAS

We ran GWAS using a three way-pipeline first used in Belinchon-Moreno et al. ^[27]^ and summarized in Fig. 1B. The first way consisted of a conventional GWAS using genome-wide SNPs. The second way used variant nodes from a pan-NLRome graph built from 143 melon lines. The third way used k-mers extracted from 21 NLR-rich sequences that served to build the pan-NLRome. As an additional analysis, we performed a pseudo-GWAS using non-redundant annotated *Vat* alleles as variants.

We used the multi-locus mixed model (MLMM) implemented in the R package mlmm ^[51]^ to perform all GWAS. For SNP-based GWAS, we ran a multistep approach, including each top SNP from previous steps as cofactors in the model until no further SNPs were found below the defined threshold or no more phenotypic variance was under genetic control. We compared four models: a baseline model (no correction), a kinship-only model (K), a model accounting for population structure (Q5, five inferred subpopulations), and a combined model including both kinship and structure (KQ5). Model selection was guided by QQ plots generated with the qqman R package ^[52]^. For GWAS restricted to the pan-NLRome, whether based on node or k-mer variants, we performed only the first step of MLMM and evaluated baseline, K, Q5 and KQ5 models. For allele-based GWAS, we just applied the simple model without correction, as this correction was already done in the previous GWAS that identified the *Vat* genes as candidates and then a double correction would generate false negatives as some *Vat* alles are restricted to a single structure group. To avoid false positives inherent to a model without correction, we performed a haplotype analysis (see “Classification of *Vat* homologs based on response to aphid-transmitted virus” section).

We used Bonferroni-corrected significance thresholds with a genome-wide error risk of α = 0.05 corrected by the number of independent variants. As presented in Belinchon-Moreno et al. ^[27]^, the number of independent tests was estimated using Gao’s method ^[53]^ with a 100 kb sliding window for SNP-based GWAS. For k-mer GWAS, independence was approximated by counting distinct P/A patterns across accessions, since k-mers linked to the same variant are assumed share the same pattern. For graph-based GWAS, the number of independent nodes was defined from top-level snarls. For allele-based GWAS, all *Vat* alleles were treated as independent.

We applied all GWAS and post-GWAS (presented hereafter) using the MultiVarGWAS Snakemake workflow (https://forge.inrae.fr/gafl/users/javier_belinchon/MultiVarGWAS).

### Post-GWAS analyses

#### Local linkage disequilibrium for QTL identification after GWAS

We delineated QTLs from SNP-based GWAS results by examining local linkage disequilibrium (LD) around the top-significant SNPs. For each top SNP, we calculated pairwise LD (r²) within a 10 Mb window using the Measure.R2VS function from the LDcorSV R package ^[54]^, accounting for both population structure and kinship. QTL boundaries were then determined using a sliding window approach: starting from the top SNP, the window was iteratively expanded 100 kb left and right until no SNPs exceeded an r² threshold of 0.2 within that window. We repeated this process for subsequent significant SNPs located outside previously defined QTLs until all significant SNPs were encompassed within defined QTL regions.

#### Candidate gene identification within QTLs, haplotype analysis and local kinship estimation

We retrieved genes located within the identified QTL regions from the gene annotation of ANSO-77 ^[37]^. For each identified gene within a given QTL, we examined whether SNPs were present in coding or non-coding regions and calculated the pairwise r^2^ between each SNP and the top-associated SNP of the QTL. We considered a gene as candidate if it carried at least one SNP in strong linkage disequilibrium with the top-significant SNP, and if the SNP was located within a coding region.

We computed local kinship matrices for each QTL using only the SNPs within the corresponding region from the GWAS SNP dataset and applying the function A.mat() from the R package rrBLUP. We clustered local kinship matrices using hierarchical clustering to identify patterns of relatedness across accessions. We assigned haplotypes by grouping accessions with similar local kinship profiles, allowing the pooling of resistant and susceptible lines according to their potential source of resistance.

#### Anchoring of top-associated k-mers and nodes

We mapped significantly-associated k-mers and nodes highlighted by targeted GWAS using the procedures detailed in Belinchon-Moreno et al. ^[27]^. To map top-significant k-mers, we first identified the accessions containing them. K-mers with identical p-values, typically tagging genomic regions in complete LD, were grouped and locally assembled with Minia v. 3.2.6, with parameters ‘”kmer-size 15 -abundance-min 1 -no-tip-removal”. We aligned the resulting k-mer contigs to the NLR assemblies containing them using BLAST v. 2.12, adding parameters “-word_size 7 -outfmt “6 std qlen” -max_target_seqs 5”, and we retained hits with 100% identity. Finally, we added gene annotations on the genomic regions with mapped k-mers using bedtools v. 2.29.2 ^[55]^. When mapping k-mers to the whole-genome assembly of ANSO-77, we retained the match with the lowest e-value for each k-mer, even when sequence identity was below 100%. This approach allowed us to retain information for all k-mers when using a single reference genome.

To anchor top-associated nodes on linear assemblies, we first determined their positions along accession-specific pan-NLRome graph paths using the odgi position command ^[56]^ included in PGGB v. 0.5.1. We then extracted the corresponding graph paths in FASTA format with vg paths from vg v. 1.63.1. Because the sequences used to construct the pan-NLRome graph were trimmed at their dangling ends compared to the ANSO-77 reference, the coordinates of nodes along the resulting graph paths did not necessarily correspond directly to their coordinates on the original annotated sequences. To retrieve the positions of top-associated nodes on original assemblies, we lifted the coordinates on the graph paths to the original annotated sequences used for graph construction, retrieved from Belinchon-Moreno et al. ^[27]^, using Minimap2 v. 2.24 ^[57]^ with parameter –f 0.02.

#### Classification of Vat homologs based on response to aphid-transmitted virus

We classified *Vat* homologs with four R65aa motifs ^[28]^ (*Vat* 4R65aa) into susceptible, intermediate-resistant, and resistant groups, based on the response of accessions carrying these homologs to CMV transmitted by five different clones of *A. gossypii* (C6, C9, CUC1, GWD2 and NM1). We selected phenotypes for 31 accessions carrying a single *Vat* homolog with four R65aa motifs, and four accessions carrying two different *Vat* 4R65aa homologs. To assign *Vat* homologs to susceptible/resistant groups in response to each aphid clone, we selected accessions carrying each *Vat* 4R65aa homolog along with the controls (VEDRANTAIS and MARGOT) phenotyped in the same incomplete blocks. We summarized the total number of plantlets presenting or lacking CMV symptoms within each group (accessions tested, resistant control, and susceptible control). For each *Vat* homolog x aphid clone combination, we performed chi-squared tests to determine whether there were significant differences between group pairs (resistant control vs susceptible control, resistant control vs tested accessions, susceptible control vs tested accessions). As some group pairs presented small or uneven sample sizes, we used Chi-squared tests with p-value estimation by Monte Carlo simulation. Simulated p-values were computed using one million permutations, providing a robust estimation of significance. All chi-squared analyses were performed using the R base function chisq.test() with B = 1,000,000 and adding the simulate.p.value = TRUE option. We assigned four levels of significance based on obtained p-values as follows: p-value ≤ 0.01, ***; p-value ≤ 0.01, **; p-value ≤ 0.05, *; p-value > 0.05, ns. We compared the significance levels of each *Vat* homolog against both resistant and susceptible control groups, accounting for the significance level observed between the two controls themselves. If the significance level comparing accessions carrying a given *Vat* homolog to the resistant control was greater or equal than that between resistant and susceptible controls, and the significance level compared to the susceptible control was lower than between controls, the homolog was classified as susceptible. Conversely, if the significance level compared to the susceptible control was equal to or higher than that between controls, and lower when compared to the resistant control, the homolog was classified as resistant. If the significance level against both resistant and susceptible controls was equal to or higher than that observed between the controls, the homolog was classified as intermediate-resistant.

#### Phylogenetic tree generation

We retrieved *Vat* homolog proteins with four R65aa motifs ^[28]^ from Belinchon-Moreno et al. ^[27]^ and we performed phylogenetic analyses to assess their genetic relatedness. We performed protein alignments using MAFFT v. 7.490 ^[58]^ with default parameters. We input these alignments to IQ-TREE v. 1.6.12 ^[59]^, adding parameters ‘-bb 10,000 -alrt 10,000’, in order to generate Maximum Likelihood phylogenetic trees. ModelFinder, which is embedded in IQ-TREE, was used to find the best-fit substitution model. The model selected was HIVb+F+I. We displayed and annotated the phylogenetic tree generated with IQ-tree using iTOL v. 7 ^[60]^.

## Results

### Bulk Segregant Analysis confirmed that resistance from ANSO-77 to the CUC1 clone of *A. gossypii* maps outside the *Vat* region

Resistance to *A. gossypii* in melon has been associated with the *Vat* cluster on chromosome 5 ^[23,28]^, as well as with several other independent loci ^[61]^. However, the genetic control of resistance to the CUC1 clone remains unknown. To determine whether the partial resistance of ANSO-77 to CUC1 colonization was associated with the *Vat* cluster, we first generated an F_2_ population from a cross between VEDRANTAIS (susceptible to colonization by the *A. gossypii* CUC1 clone) and ANSO-77 (partially resistant). Using genetic markers targeting the *Vat* genomic region, we identified 38 F_2_ plants carrying the *Vat* cluster from VEDRANTAIS, and 49 F_2_ plants carrying the *Vat* cluster from ANSO-77. Phenotyping of these two groups for aphid acceptance, defined as the initial settling of aphids, and colonization, defined as their ability to establish on the plant, revealed no significant differences, evidencing that the resistance of ANSO-77 to CUC1 colonization was not associated with its *Vat* cluster (Fig. S3).

Based on this result, we generated a second F_2_ population from a cross between ANSO-77 and DOUBLON (highly susceptible to CUC1 colonization) to identify resistance loci segregating independently of the *Vat* cluster. The resulting 576 F_3_ families, derived from selfing individual F_2_ plants, were initially phenotyped using a mass inoculation approach, which allowed us to identify 202 F₃ families with extreme phenotypic scores. Individual-plant phenotyping of these selected families under controlled infestation conditions confirmed a wide range of CUC1 colonization levels. Based on a normalized colonization score relative to resistant and susceptible controls, we selected 38 F₃ families at the two extremes of this distribution to constitute resistant and susceptible bulks (Fig. S4A). The resistant bulk showed a mean aphid colonization score of 5.0, close to that of the resistant control (mean = 4.4), whereas the susceptible bulk exhibited a mean score of 7.6, similar to that of the susceptible control (mean = 7.5) (Fig. S4B).

The selected 76 F₃ families (2 x 38) were represented in resistant and susceptible DNA bulks through equimolar pooling of DNA from their corresponding F₂ parents. Whole-genome short-read sequencing yielded average sequencing depths exceeding 164× for the bulks and 21× for the ANSO-77 and DOUBLON parental lines. In all cases, more than 99% of the reads mapped to the parental reference genomes. Variant calling using the genomes of ANSO-77 (369.5 Mb, including unassembled contigs) and DOUBLON (362.6 Mb, including unassembled contigs) as references identified 665,603 and 677,435 high-confidence variants, mostly SNPs, with few INDELs shorter than 10 bp. The density plots of filtered variants are depicted in Fig. 2A and S5A. We performed QTL-seq analysis based on the calculation of a Δ(SNP-index) for both resistant and susceptible bulks. We visually identified a very large QTL on chromosome 12 (Fig. 2B and S5B), potentially containing multiple linked resistance loci, and suggesting the presence of long haplogroups. Using a 99% Takagi-based confidence interval to define QTL boundaries, we detected four distinct genomic regions forming this large QTL, spanning in total 17.47 Mb in ANSO-77 and 17.53 Mb in DOUBLON (Table S2). Excluding interspaced regions where the Δ(SNP-index) fell below the threshold, the effective QTL region covered 11.33 Mb in ANSO-77 and 12.09 Mb in DOUBLON (Table S2). The positive Δ(SNP-index) obtained when using the ANSO-77 genome assembly as the reference evidenced that the resistance underlying this QTL is contributed by ANSO-77, the resistant parent. Complementary G’ analyses also evidenced this QTL on chromosome 12 (Fig. 2C and S5C), defining an interval of 20.94 Mb in ANSO-77 and 22.28 Mb in DOUBLON (Table S2). In addition, G’ analysis identified a second putative QTL spanning 413.24 Kb and located on the unassembled contigs of the ANSO-77 genome (Fig. 2C).

**Fig. 2.**
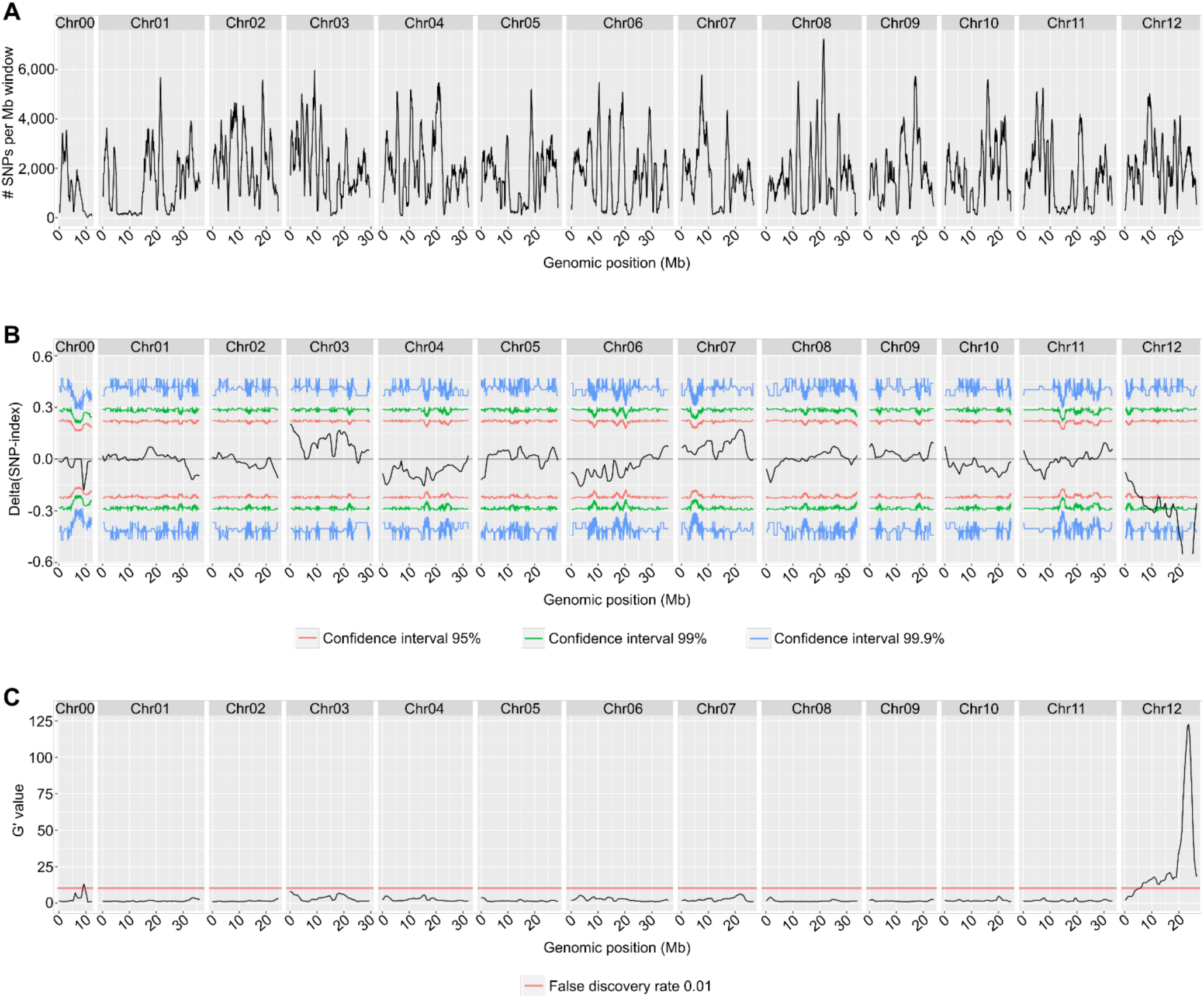
QTLs associated with resistance to A. gossypii clone CUC1 colonization, identified by QTL-seq using the ANSO-77 genome as reference. All analyses were performed using a 1 Mb sliding window. **A,** Distribution of SNPs and short INDELs within each 1 Mb smoothing window along the 12 ANSO-77 chromosomes and unassembled contigs. **B,** Tricube-smoothed Δ(SNP-index) distribution along the 12 ANSO-77 chromosomes and unassembled contigs. Red, green and blue lines indicate the 95%, 99% and 99.9% two-sided confidence intervals, respectively. **C,** Tricube-smoothed G’ values distribution along the 12 ANSO-77 chromosomes and unassembled contigs. The red horizontal line represents a genome-wide false discovery rate threshold of 0.01.

Together, these results identified a large genomic region spanning almost half of chromosome 12 in the ANSO-77 genome underlying resistance to the *A. gossypii* CUC1 clone. To identify resistance loci beyond those segregating in this biparental cross and to assess the contribution of natural genetic variation, we next extended the analysis to a genetically diverse panel of melon accessions using GWAS.

### GWAS identified several QTLs of response to *A. gossypii* CUC1 clone

We assembled a panel of 212 melon accessions (PA_colonization_ panel) to identify additional genomic regions associated with variation in response to the *A. gossypii* CUC1 clone and evaluated their phenotypic responses. Plant acceptance, assessed 72 h after placing ten aphids on each plant, ranged from 2.1 to 9.1 (mean = 6.5) aphids remaining on the plant. In addition to the resistant control VIRGOS, five accessions (SLK-V-052, SMOOTH CALCUTTA, PASTIS_2, CHINA_51, and SAN ILDEFONSO) showed low acceptance by aphids, with fewer than three of the ten aphids remaining on the plant after three days. Most of these resistant accessions originated from India or the Far East (Table S1). Aphid colonization, observed after seven days, ranged from 2.8 to 10.8 (mean = 8.1), with the least colonized accessions originating from highly diverse geographical regions. Broad-sense heritability estimates were 79.7% and 72.9% for acceptance and colonization, validating the experimental protocols. We estimated BLUPs using LMMs to predict phenotypic values for each accession (Table S3). For acceptance, we included the test as a random effect. For colonization, test was fitted as a random effect and acceptance as a fixed effect, allowing the extraction of a derived phenotype, termed multiplication, which reflects a different step of the aphid/melon interaction than that measured by acceptance. Acceptance and multiplication BLUPs were partially correlated (r² = 0.71) (Fig. S6), suggesting partially shared genetic control of resistance to aphid establishment and multiplication.

We estimated the genetic structure of the 212 accessions included in the PA_colonization_ panel using the STRUCTURE software (Table S4). We inferred five groups (Q5) as the most relevant genetic organization (Fig. S7A,B), closely reflecting the known evolutionary history of botanical groups and geographical origins. The distribution of acceptance and multiplication BLUPs across structure groups revealed both phenotypic variation within groups and phenotypic differences among groups. Particularly, the genetic group 5.3, mainly composed of accessions from the Far East belonging to botanical groups *conomon*, *makuwa* and *chinensis*, showed the lowest average acceptance and multiplication (Fig. S7E,F). In contrast, accessions from the genetic group 5.5 (primarily originating from Europe, the Middle East, Central Asia, or North America), as well as those reflecting admixture, displayed the highest averages (Fig. S7E,F).

We applied a multi-locus SNP-based GWAS model using the MA_colonization_ matrix (713,485 evenly distributed SNPs across the ANSO-77 genome). Among the four tested GWAS models (null, K, Q5 and KQ5), the MLMM model incorporating both kinship and five genetic structure groups (KQ5) yielded p-values that produced Q-Q plots most closely following the bisector, regardless of the phenotype (acceptance or multiplication) (Fig. S8A,B). Using this model, we identified SNPs over the Bonferroni-corrected threshold on chromosomes 3, 8 and 12 for acceptance, and on chromosomes 5 and 12 for multiplication (Fig. 3A). Local LD calculation around the top-significant SNPs identified three QTLs for aphid acceptance and six QTLs for aphid multiplication. Detailed information for each detected QTL is available in Table S2. Analysis of variance (ANOVA) of the phenotypic distributions across the three genotype classes of the top SNPs revealed a significant effect (p < 0.001) for all QTLs (Fig. 3B).

**Fig. 3.**
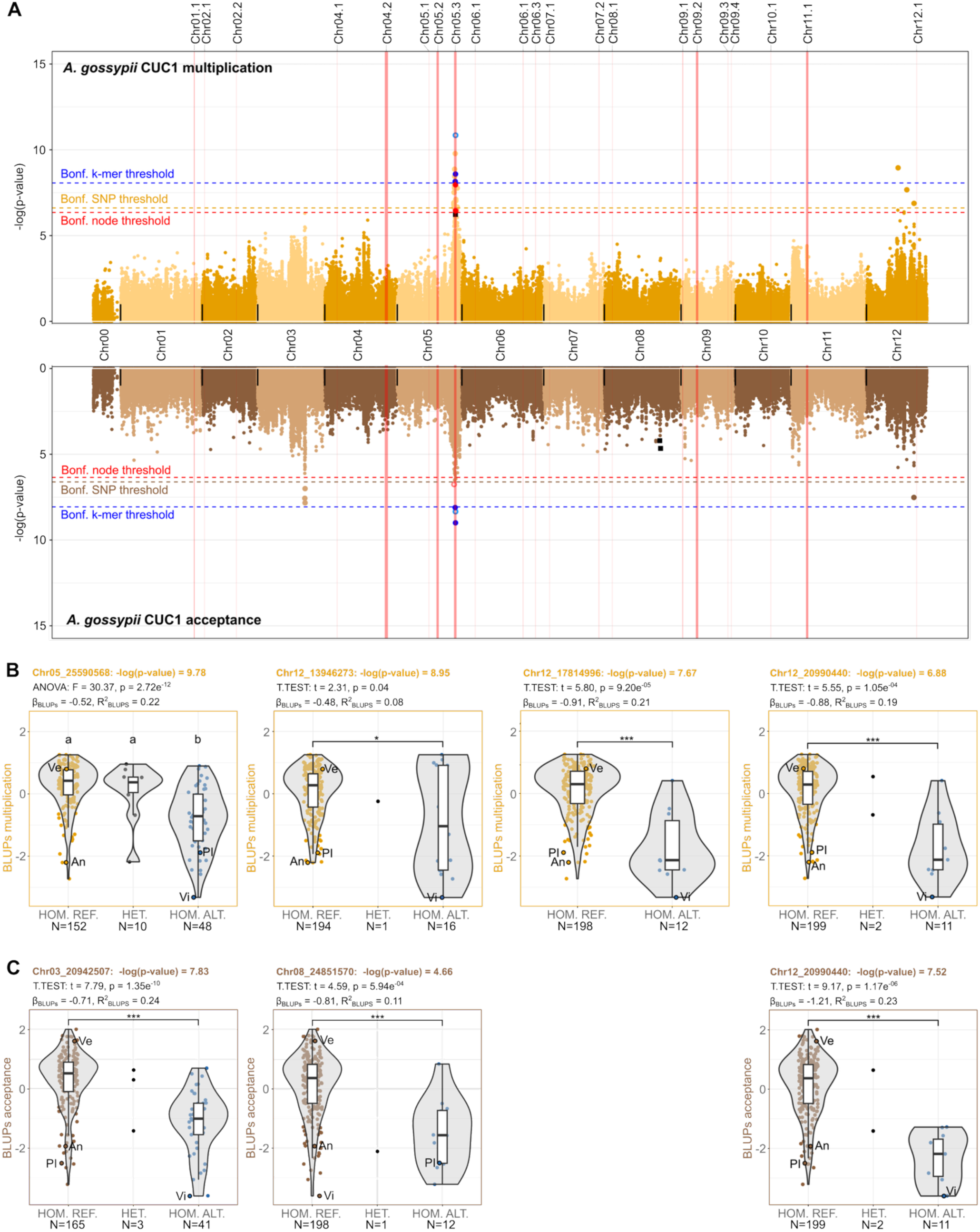
QTLs associated with plant responses to *A. gossypii* clone CUC1 multiplication and acceptance, identified by GWAS. **A,** Miami plot showing mirrored Manhattan plots for multiplication (top) and acceptance (bottom). SNP-based GWAS was performed at the whole-genome scale, enabling the screening of all chromosomes, whereas pan-NLRome graph node- and k-mer-based GWAS were restricted to 21 targeted genomic regions containing NLRs, providing a higher-resolution analysis of these regions. All p-values on the Manhattan plot correspond to the first MLMM step. Whole-genome SNPs are shown in orange (multiplication) and brown (acceptance); SNPs retained in the second step of MLMM are highlighted as black squares. Significant k-mers and pan-NLRome graph nodes are shown with filled dots (blue for k-mers and red for nodes) when their sequences matched the ANSO-77 genome at 100% identity, and with lighter, unfilled symbols when they did not. Significant k-mers lacking a 100% identity match to ANSO-77 were positioned at the genomic position with the highest sequence similarity, whereas absent graph nodes lacking the ANSO-77 sequence path were positioned at the genomic position of the closest graph node containing the ANSO-77 genome sequence. Black vertical lines delimit chromosomes. Vertical red bars locate NLR-containing genomic regions, scaled to region size. Horizontal dashed lines indicate Bonferroni-corrected thresholds (α = 0.05, adjusted for the estimated number of independent variants). **B and C,** Distribution of BLUPs according to allelic configuration at the top-significant SNPs for each reported QTL, based on multiplication (B) and acceptance (C). HOM. REF: homozygous reference; HET: heterozygous; HOM. ALT: homozygous alternative. Dots represent individual accessions. Ve: VEDRANTAIS (S); An: ANSO-77 (R); PI: PI 414723 (R); Vi: Virgos (R). N indicates the number of accessions per allelic configuration. Only groups of at least five accessions were considered. Group differences were tested using t-tests (two groups) or ANOVA followed by Tukey’s HSD (≥ 3 groups). T-test significance levels are indicated as * (p < 0.05), ** (p < 0.01), and *** (p < 0.001). Linear regression coefficients (β_BLUPs_) represent allelic effects, and R² values indicate the proportion of phenotypic variance explained.

**The QTL on chromosome 3**, associated with variation in acceptance, spanned 969.06 kb, contained three significant SNPs (Table S2), and encompassed 86 annotated genes (Fig. S9). Among candidate genes, Chr03.1000694, encoding a MYB33 transcription factor, and Chr03.1000687, encoding an ethylene insensitive 3-like 1 (EIL1) gene, stood out due to their well-established role in aphid resistance across multiple species ^[4–8]^. These two genes, closely linked on chromosome 3, may act in concert to regulate phloem-based defenses through coordinated crosstalk between MYB-mediated transcriptional control of ethylene signaling via EIL1, similar to the mechanisms described in other plant species ^[8,9]^. **The strongly supported SNP peak on chromosome 5** (multiplication) overlapped the largest NLR cluster in melon ^[62]^, which contains the well-characterized *Vat* resistance cluster. The most significant SNP located 51.7 kb from the M5 marker ^[28]^ associated to the *Vat* cluster. Local LD analysis partitioned this peak into three closely located QTLs, the second of which spanned 543.56 kb and fully encompassing the *Vat* M5-M4 genomic region ^[28]^ (Table S2). This QTL is analyzed in detail in the next section. **The QTL on chromosome 8** (acceptance) was detected in the second MLMM step, after incorporating the top SNP Chr03_20942507 as a fixed effect (Fig. 3A and Table S2). This 490.51 kb region contained 26 genes, of which three were highlighted as candidate genes. However, none of them showed features suggesting a role in resistance to biotic stresses. **Three QTLs on chromosome 12** (two associated with variation in multiplication, and the third common to multiplication and acceptance, Table S2) were weakly supported, each being represented by a single SNP exceeding the corrected Bonferroni threshold (Fig. 3A). Nevertheless, phenotypic differences between reference and alternative alleles at these SNPs were statistically significant in all cases (Fig. 3B,C). Notably, all three QTLs detected on chromosome 12 by GWAS co-localized within the broader genomic region previously identified by BSA-NGS using the colonization phenotype, which integrates acceptance and multiplication. Although BSA-NGS analysis indicated that resistance in this genomic region originates from ANSO-77, this accession carried the allele of susceptibility at the three GWAS-identified QTLs (Fig. 3B,C). The allele of resistance contributed by ANSO-77 was likely not detected by GWAS due to its low frequency in the diversity panel. Indeed, SNP-based haplotype analysis around the BSA-NGS peak showed that ANSO-77 harbors a unique haplotype at this locus.

Having characterized the genetic architecture of melon responses to CUC1 acceptance and multiplication, we next expanded the GWAS analysis to aphid attractiveness, a first layer of plant resistance acting during host selection. This trait was evaluated in 182 accessions using multiple-choice assays with the *A. gossypii* CUC1 clone; eight accessions were subsequently excluded from the GWAS because of insufficient genotyping coverage. The estimated attractiveness values ranged from 0 to 0,024 (Table S3 and Fig. S6) and were moderately correlated with aphid acceptance (r² = 0.53) and aphid multiplication (r² = 0.44) (Fig. S6). The least attractive accession was by far the Indian line PI 414723, whereas the most attractive were “Prescott à fond Blanc paris” and “Tibish Djebel Kordofan 4”, originating from France and Soudan. Genetic structure analysis on the panel of 174 accessions (MA_attractiveness_) suggested three genetic groups (Q3) as the most relevant genetic organization (Table S5 and Fig. S7C,D). The distribution of the attractiveness phenotype (normalized using a logarithmic transformation) across the genetic groups revealed large phenotypic variations within groups (Fig. S7G). The genetic group A3.2, mainly comprising accessions from the Far East belonging to botanical groups *conomon*, *makuwa* and *chinensis*, presented in average the lowest normalized attractiveness. In contrast, the genetic group A3.3, mainly composed of accessions from Europe, North America, Central Asia, and Middle East, presented the highest values. Actually, Northern American lines presented an intermediate attractiveness to the CUC1 aphid clone (0.006 in average for 16 accessions) (Table S3).

We performed SNP-based GWAS via MLMM on the normalized attractiveness phenotype, using a SNP matrix containing 533,109 SNPs evenly distributed across the genome of ANSO-77. Among the four tested GWAS models (null, K, Q3 and KQ3), Q3 presented the best fit based on Q-Q exploration (Fig. S8C). Applying this model, we identified two SNPs on chromosome 6 over the defined Bonferroni-corrected threshold (Fig. 4A and Table S2). The top SNP presented a strong additive behavior, with an allelic effect, β, of −0.52 (Fig. 4B). Local LD calculation around this top-significant SNP provided a QTL of 373.56 kb, including 21 annotated genes in ANSO-77. None of these genes emerged as strong candidates based on their predicted functions.

**Fig. 4.**
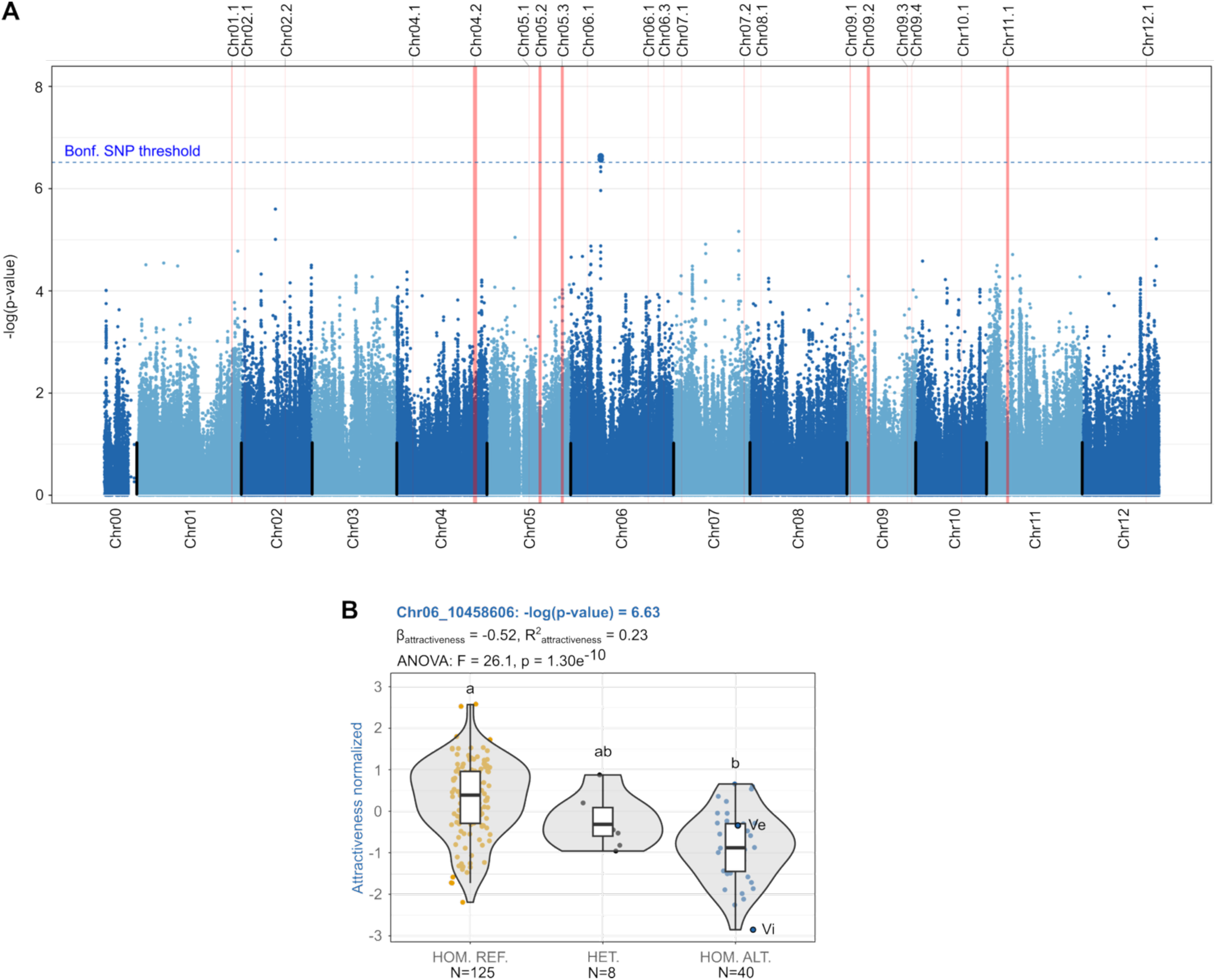
QTL associated with plant attractiveness to *A. gossypii* clone CUC1, identified by GWAS. **A,** Manhattan plot of whole-genome SNP-based GWAS results. P-values correspond to the first MLMM step. The horizontal dashed line indicates the Bonferroni-corrected threshold (α = 0.05, adjusted for the estimated number of independent variants). Vertical red bars locate NLR-containing genomic regions, scaled to their size. **B,** Distribution of normalized attractiveness phenotype according to allelic configuration at the top-significant SNP on chromosome 6. HOM. REF: homozygous reference; HET: heterozygous; HOM. ALT: homozygous alternative. Dots represent individual accessions. Ve: VEDRANTAIS (S); Vi: Virgos (R). N indicates the number of phenotyped accessions contributing to each genetic configuration. Differences among groups were tested using ANOVA followed by Tukey’s HSD. Linear regression coefficients (β_BLUPs_) indicate allelic effects, and R² values indicate the proportion of phenotypic variance explained.

Altogether, whole-genome SNP-based GWAS identified five distinct genomic regions on chromosomes 3, 5, 6, 8 and 12 associated with variation in three traits capturing different stages of the melon-aphid (CUC1 clone) interaction. Among these, the chromosome 5 signal associated with resistance to multiplication was the only overlapping an NLR-containing region, corresponding to the well-characterized *Vat* locus. We therefore focused subsequent high-resolution analyses on this region using k-mer- and graph-based targeted GWAS.

### Converging evidence implicated *Vat* alleles with four R65aa motifs in resistance to CUC1 *A. gossypii* clone

To further investigate the role of the *Vat* NLR cluster in resistance to multiplication by the CUC1 clone of *A. gossypii*, we applied targeted k-mer- and graph-based GWAS following the methodology described in Belinchon-Moreno et al. ^[27]^. These approaches complemented the whole-genome SNP analysis by capturing sequence variation, from SNPs to structural variants, within 21 NLR-containing regions without restricting variant discovery to sequences anchored to a single reference genome. The genomic locations of these targeted NLR regions are indicated by red vertical bars in the Manhattan plots (Fig. 3A). We retained 132 accessions with available CUC1 multiplication phenotypes and with their NLRome sequenced with long reads (forming part of the PA_NAS_ panel). These phenotypes were linked to the presence/absence of 31-mers within NLR regions (k-mer-based GWAS) or to the presence/absence of the sequence path of each accession on graph nodes (graph-based GWAS). We applied MAF cutoffs of 0.06 to the P/A matrices of k-mers and graph nodes, retaining ∼17.05 million k-mers and 330,579 nodes for GWAS. Among the tested targeted GWAS models (K, Q5 and KQ5), KQ5 consistently presented the best fit based on Q-Q exploration (Fig. S8D-G) and was thus selected for further analyses.

K-mer-based GWAS identified eight significant k-mers. Local assembly of k-mers sharing identical p-values resolved these signals into three k-mer blocks that aligned to the QTL on chromosome 5 already identified by SNP-based GWAS (Fig. 3A). Detailed analysis on three representative accessions with extreme phenotypes (ANSO-77, VEDRANTAIS and PI 414723) showed that the two most significant k-mer blocks located within the *Vat* M5-M4 region and aligned to intergenic sequences in all the assemblies (Fig. 5). The third-significant k-mer block mapped ∼53 kb upstream of M5. Graph-based GWAS detected three significant nodes, all corresponding to variants within the *Vat* M5–M4 region and also mapping to intergenic regions (Fig. 5). The second-most significant k-mer and the first and third top-significant nodes showed general positive allelic effects on resistance to CUC1 multiplication (Fig. S10). These markers were located near *Vat* alleles carrying 4R65aa and 5R65aa motifs within their LRR domains (Fig. 5). Overall, targeted k-mer- and graph-based GWAS increased the resolution of the chromosome 5 association signal previously detected with SNP-based GWAS and provided converging evidence that resistance to CUC1 multiplication is likely encoded within the M5–M4 *Vat* region. However, these approaches did not allow the identification of specific *Vat* alleles underlying this resistance.

**Fig. 5.**
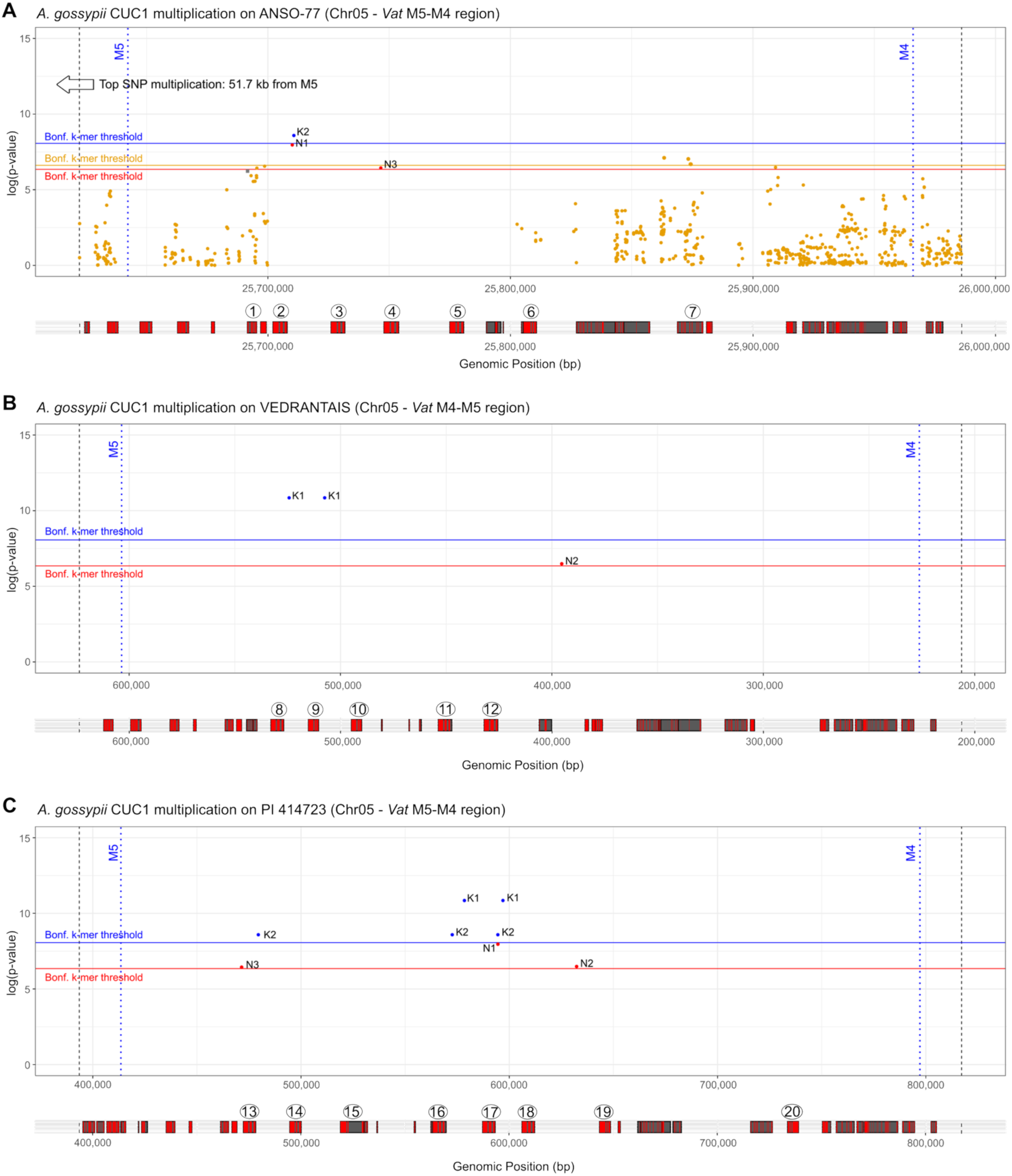
SNP-, k-mer and graph-based GWAS results for resistance to multiplication by the CUC1 clone of *A. gossypii*, focusing on the *Vat* genomic region defined by the M5 and M4 markers. **A,** Significant SNPs, k-mers and graph nodes on ANSO-77, resistant to *A. gossypii* CUC1 multiplication but carrying a *Vat* region not associated with resistance. **B,** Significant k-mers and graph-nodes present in VEDRANTAIS, a susceptible accession. **C,** Significant k-mers and graph-nodes present in PI 414723, a resistant accession. Blue dashed vertical lines represent the positions of M5 and M4 markers. Black dashed vertical lines are located 20 kb upstream and downstream of M5 and M4, respectively. Horizontal colored lines indicate the Bonferroni-corrected significance thresholds, calculated with α = 0.05, adjusted for the estimated number of independent variants. Annotated genes within the regions are plotted below the corresponding Manhattan plots. Red blocks represent exons, and black blocks represent introns. Gene IDs are as follows: 1, ME00509#1#utg45_Chr05.3_000100; 2, ME00509#1#utg45_Chr05.3_Vat1_4R65aa; *3, ME00509#1#utg45_Chr05.3_Vat2_3R65aa; 4, ME00509#1#utg45_Chr05.3_Vat3_5R65aa; 5, ME00509#1#utg45_Chr05.3_Vat4_3R65aa; 6, ME00509#1#utg45_Chr05.3_Vat5_1R65aa; 7, ME00509#1#utg45_Chr05.3_000115; 8, ME01000#1#utg194_Chr05.3_Vat1_5R65aa; 9, ME01000#1#utg194_Chr05.3_Vat2_2R65aa; 10, ME01000#1#utg194_Chr05.3_Vat3_2R65aa; 11, ME01000#1#utg194_Chr05.3_Vat4_4R65aa; 12, ME01000#1#utg194_Chr05.3_Vat5_7R65aa; 13, ME00594#1#utg4_Chr05.3_Vat1-4R65aa; 14, ME00594#1#utg4_Chr05.3_Vat2_2R65aa; 15, ME00594#1#utg4_Chr05.3_Vat3_1R65aa, 16, ME00594#1#utg4_Chr05.3_Vat4-4R65aa; 17, ME00594#1#utg4_Chr05.3_Vat5-5R65aa; 18, ME00594#1#utg4_Chr05.3_Vat6-5R65aa; 19, ME00594#1#utg4_Chr05.3_Vat7-1R65aa; 20, ME00594#1#utg4_Chr05.3_Vatrev*.

To further investigate resistance carried by the *Vat* cluster at the allele level, we recovered manual annotation of *Vat* alleles across 143 accessions from Belinchon-Moreno et al. ^[27]^ and clustered alleles sharing identical amino acid sequences. This yielded a P/A matrix of 167 non-redundant *Vat* alleles across 132 accessions phenotyped for CUC1 multiplication. Applying a MAF > 0.03 (allele present in at least four accessions) retained 32 alleles for further analysis. We performed an ‘Allele Association Study’ with this matrix using the first step of MLMM under a null model to screen individual *Vat* alleles for association with the resistance phenotype. Population-structure and kinship corrections were not applied here as these corrections were already incorporated in k-mer- and graph-based GWAS, which identified the *Vat* region as being associated to resistance to CUC1 multiplication. Applying the same correction again could mask biologically relevant associations, given the strong structuring of some *Vat* alleles across genetic groups. Six alleles showed a positive effect (1R65aa_B, 1R65aa_E, 4R65aa_B, 4R65aa_C and VatRev_B and 2R65aa_C) and four presented negative effect (2R65aa_A, 4R65aa_A, 5R65aa_A and 7R65aa_A) on resistance to CUC1 multiplication (Fig. 6A). Haplotype analyses further highlighted 4R65aa_B and 4R65aa_C as the strongest candidates for resistance to CUC1 multiplication (Fig. 6B). Indeed, allele 1R65aa_E was consistently associated with 4R65aa_B and 4R65aa_C, with ME01367 being the only exception and showing a susceptible phenotype. Similarly, VatRev_B was always linked to 4R65aa_B or 4R65aa_C. Alleles 1R65aa_B and 2R65aa_C occurred both in association with 4R65aa_B or 4R65aa_C and independently; however, haplotype analyses indicated no evidence of resistance when they occurred in the absence of either 4R65aa_B or 4R65aa_C.

**Fig. 6.**
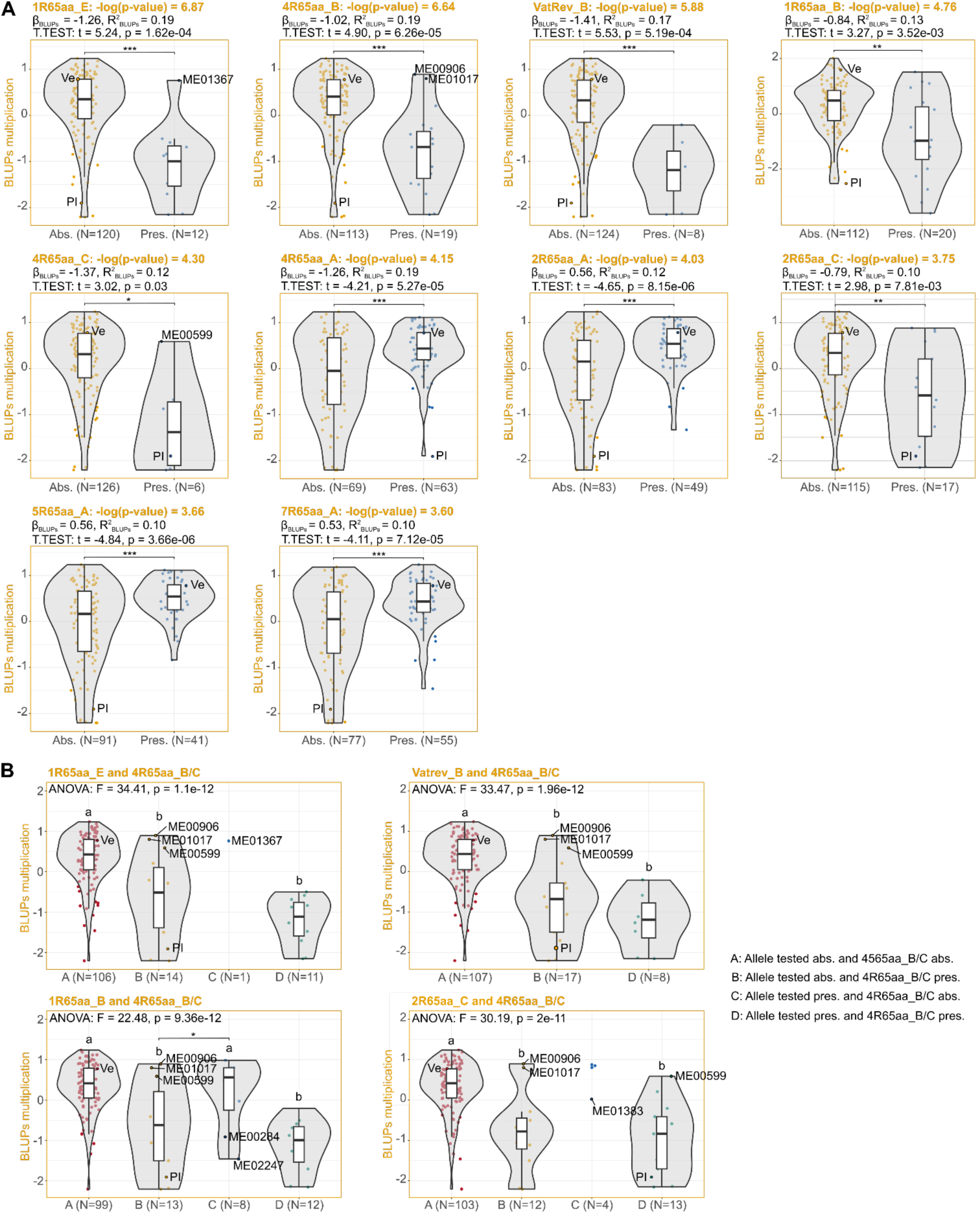
Candidate *Vat* homologs associated with plant responses to *A. gossypii* clone CUC1 multiplication, identified by allele-based GWAS. **A,** Distribution of BLUPs across accessions carrying (pres.) or lacking (abs.) each reported *Vat* homolog. Dots represent individual accessions. Ve: VEDRANTAIS (S); Vi: Virgos (R). N indicates the number of accessions per group. Differences between groups were tested using t-tests: * (p < 0.05), ** (p < 0.01), and *** (p < 0.001). Linear regression coefficients (β_BLUPs_) indicate the effect of the presence of each *Vat* homolog, and R² values indicate the proportion of phenotypic variance explained. **B,** Distribution of BLUPs across the four possible haplotypes defined by the presence/absence of 4R65aa_B and 4R65aa_C combined with the rest of homologs positively associated with the reduced aphid multiplication. Only groups with ≥ 5 accessions were considered. Differences between groups were estimated using ANOVA followed by Tukey’s HSD post-hoc. Significance between B (allele tested absent and 4R65aa_B or 4R65aa_C present) and C (allele tested present and 4R65aa_B or 4R65aa_C absent) groups are indicated as * (p < 0.05), ** (p < 0.01), and *** (p < 0.001). ME00284 carries a 4R65aa_Q homolog. ME00247 carries 4R65aa_G homolog. ME01383 carries a 4R65aa_N homolog.

To avoid false positives inherent to a model without correction and further validate these findings, we performed an additional SNP-based GWAS after excluding accessions carrying 4R65aa_B, 4R65aa_C and eight additional *Vat* homologs containing four R65aa motifs that were subsequently linked to CUC1 resistance (4R65aa_E, 4R65aa_F, 4R65aa_G, 4R65aa_H, 4R65aa_M, 4R65aa_N, 4R65aa_K, 4R65aa_I, see next paragraph). The GWAS on the remaining 98 accessions revealed no association peak on chromosome 5 (Figs. S8D and S11), consistent with the hypothesis that resistance to CUC1 multiplication is primarily driven by these *Vat* homologs with four R65aa motifs.

### Resistance assays with aphid-inoculated virus revealed resistant and susceptible *Vat* alleles against five *A. gossypii* clones

The *Vat* 4R65aa_B allele has previously been shown to confer resistance to CMV when inoculated by certain clones of *A. gossypii* ^[28]^. This resistance is likely triggered by an effector present in the aphid saliva, *a priori*, specific to each aphid clone ^[18]^. The GWAS presented above further highlighted the relatively frequent *Vat* alleles 4R65aa_B and 4R65aa_C as likely candidates for resistance to CUC1 multiplication. This prompted us to investigate the resistance spectrum of *Vat* 4R65aa homologs across different *A. gossypii* clones. To this end, we used the virus-transmission resistance phenotype that has the advantage of not being confounded by the effect of QTLs likely linked to aphid acceptance/multiplication on chromosomes 3, 8 and 12.

We considered 20 *Vat* 4R65aa homologs identified in the 143 accessions composing the PA_NAS_ panel, called from A to T. Among these accessions, 108 carried at least one *Vat* 4R65aa homolog (Table S6). The 4R65aa_A allele was present in more than half of the accessions, followed by 4R65aa_B in nearly 20%, whereas the remaining homologs occurred in 1-7 accessions each. We selected 31 accessions carrying a single *Vat* 4R65aa homolog and four accessions carrying two different *Vat* 4R65aa homologs (Table S6), resulting in 35 accessions phenotyped for resistance to CMV inoculated by five *A. gossypii* clones (C6, C9, CUC1, GWD2 and NM1). By comparing the phenotypes of the 35 accessions with those of resistant and susceptible controls within the same incomplete blocks, we classified *Vat* 4R65aa alleles as conferring resistance, intermediate resistance or susceptibility to each *A. gossypii* clone (Fig. 7A). From the 20 non-redundant *Vat* 4R65aa homologs, 14 conferred partial or complete resistance to CMV inoculated by at least one clone, 11 to at least two clones, and nine to at least three clones. Two homologs, 4R65aa_O and 4R65aa_P, were each found in a single assembled accession, always in association with homolog 4R65aa_C. Therefore, their link to resistance or susceptibility could only be assessed for clones to which 4R65aa_C was not associated with resistance, as it was the C6 clone. Remarkably, 4R65aa_O and 4R65aa_P were the alleles conferring the strongest CMV-resistance triggered by the C6 clone.

**Fig. 7.**
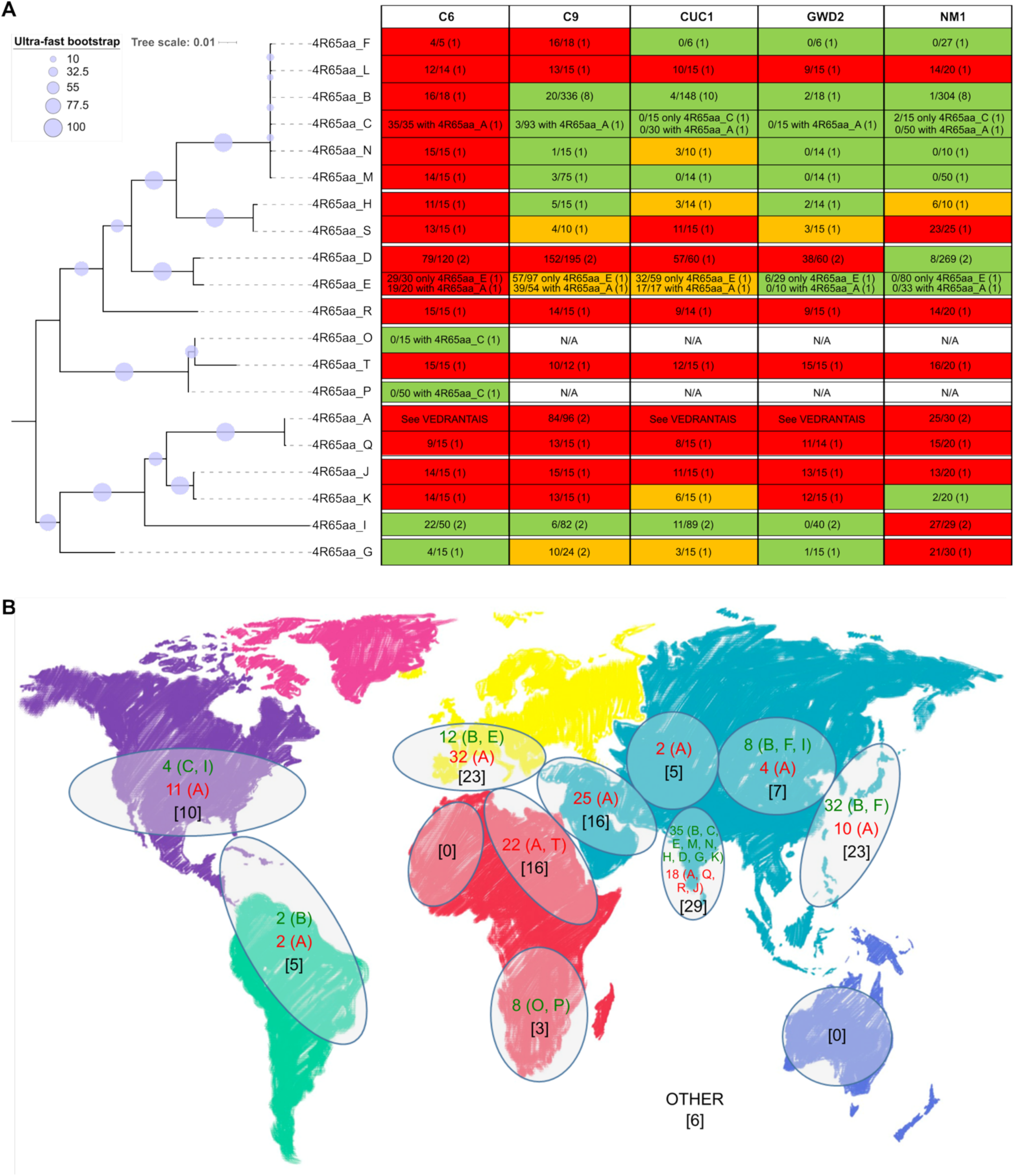
Spectrum of resistance conferred by 20 *Vat* homologs containing four R65aa motifs against five *A. gossypii* clones (C6, C9, CUC1, GWD2 and NM1). **A,** Maximum-likelihood phylogeny of 20 4R65aa *Vat* homologs (named from A to T), and their associated resistance/susceptibility patterns to the five *A. gossypii* clones. Trees were inferred with 10,000 ultra-fast bootstrap replicates and 10,000 SH-like approximate likelihood ratio tests. For each *Vat* homolog, the classification of its response to the five *A. gossypii* clones was based on chi-squared tests with one million Monte Carlo simulations, comparing the number of resistance and susceptible plantlets between accessions carrying that homolog to the resistant (MARGOT, carrying homolog 4R65aa_C) and susceptible (VEDRANTAIS, carrying homolog 4R65aa_A) controls phenotyped in the same assays. Within each cell, x/y (z) denotes the number of plantlets showing CMV symptoms (x) over the total tested (y), with z indicating the number of distinct plant genotypes considered. For clone C6, red cells indicate no significant difference from the susceptible control, and green cells indicate significant difference. For the other clones, red indicates no significant difference from the susceptible control, green no significant difference from the resistant control, and orange significant differences from both susceptible and resistant controls (significance threshold was set on the significance p-value between both susceptible and resistant controls). For 4R65aa_O and 4R65aa_P, resistance to clones C9, CUC1, GWD2 and NM1 could not be inferred because tested accessions also carried 4R65aa_C, which confers resistance to these clones. **B,** Geographic distribution of the 20 *Vat* 4R65aa alleles (named A to T) observed in 143 accessions. Numbers in brackets indicate accession counts per geographic groups; green and red values correspond to the counts of alleles of resistance and susceptibility, with allele names between parentheses.

We reconstructed the phylogeny of the 20 *Vat* 4R65aa homologs, revealing two main clades with no clear differences in resistance patterns conferred (Fig. 7A). Homologs that conferred resistance to all five tested aphid clones were observed in both clades, and each clade contained homologs associated with both narrow and broad-spectrum resistance (Fig. 7A). The first clade (4R65aa_F to 4R65aa_P) gathered 16 homologs, six of which conferred complete or partial resistance to four aphid clones, while three did not confer resistance to any of the tested clones. The second clade (4R65aa_A to 4R65aa_G) included six homologs, with two conferring complete or partial resistance to four aphid clones, while three showed no resistance to any of the tested clones (Fig. 7A). Although some groups of closely related homologs exhibited similar aphid response profiles, such as the subclade 4R65aa_B to 4R65aa_M, or the homologs pair 4R65aa_A and 4R65aa_Q, the opposite was also observed. Indeed, some closely related homologs with minimal sequence divergence exhibited markedly distinct resistance profiles. For instance, just two amino acid substitutions between 4R65aa_L to 4R65aa_F resulted in the gain of resistance to clones CUC1, GWD2 and NM1. Going back to 143 assembled accessions, those from East Asia and India carried an average of 1.3 alleles of resistance per accession, while, in all others areas, accessions carried in average less than 0.5 alleles of resistance per accession and even none in East Africa, Middle East and Central Asia (Fig. 7B). Remarkably, accessions from India carried nine different alleles of resistance with four alleles observed nowhere else, while East Asian accessions carried three different alleles of resistance with one allele observed nowhere else. This suggested that those regions have imposed highly contrasted selective pressure on the *Vat* cluster.

## Discussion

Farmers with access to aphid-resistant varieties use them extensively because they are the simplest way to control aphid infestations. The extensive cultivation of *A. gossypii*-resistant melons in France is emblematic of the success of aphid-resistance breeding of horticultural crops. MARGOT, the first aphid-resistant melon cultivar, was registered in France in the 1987 ^[24]^. Since then, the demand from European farmers has increased to the extent that all new melon cultivars recently proposed for registration are declared resistant to aphids (https://www.geves.fr/catalogue-france/).

Our understanding of plant resistance to aphids advanced significantly when plant-aphid interactions were incorporated in the fruitful ‘zig-zag’ model for plant-pathogen interactions ^[63]^. Once aphids select a plant to feed on, the molecular dialogue between the insect and the host is initiated in the plant apoplast. During this process, the aphid mouthparts (stylets) penetrate the apoplastic space and eject saliva containing pathogen-associated molecular patterns (PAMP)-like molecules. These PAMPs can be recognized by plant cell surface receptors, leading to PAMP-triggered immunity (PTI) ^[64]^ (Fig. 8A). But plants can also respond to aphids’ attack using a more specialized layer of defense, the effector-triggered immunity (ETI). For ETI, the dialogue aphid/plant is initiated within plant cells and is likely to occur via the aphid’s watery saliva, which is injected into the cytoplasm when the aphid’s stylets puncture cells on their way to reach the phloem. This watery saliva is expected to contain effectors, which may trigger ETI when recognized by the appropriate plant NLR (Fig. 8A). From a plant breeding perspective, ETI has been extensively utilized for conferring resistance to pathogens and aphids. This widespread use was likely due to the frequent alignment of ETI with the Flor’s gene-for-gene model ^[65]^, where resistance is governed by a single dominant R gene. In addition to ETI, other resistance loci, often associated with more subtle or quantitative effects, also contribute to plant resistance. Integrating them alongside major R-genes in breeding programs can help to reduce the risk of resistance breakdown and promotes more sustainable crop protection over time.

**Fig. 8.**
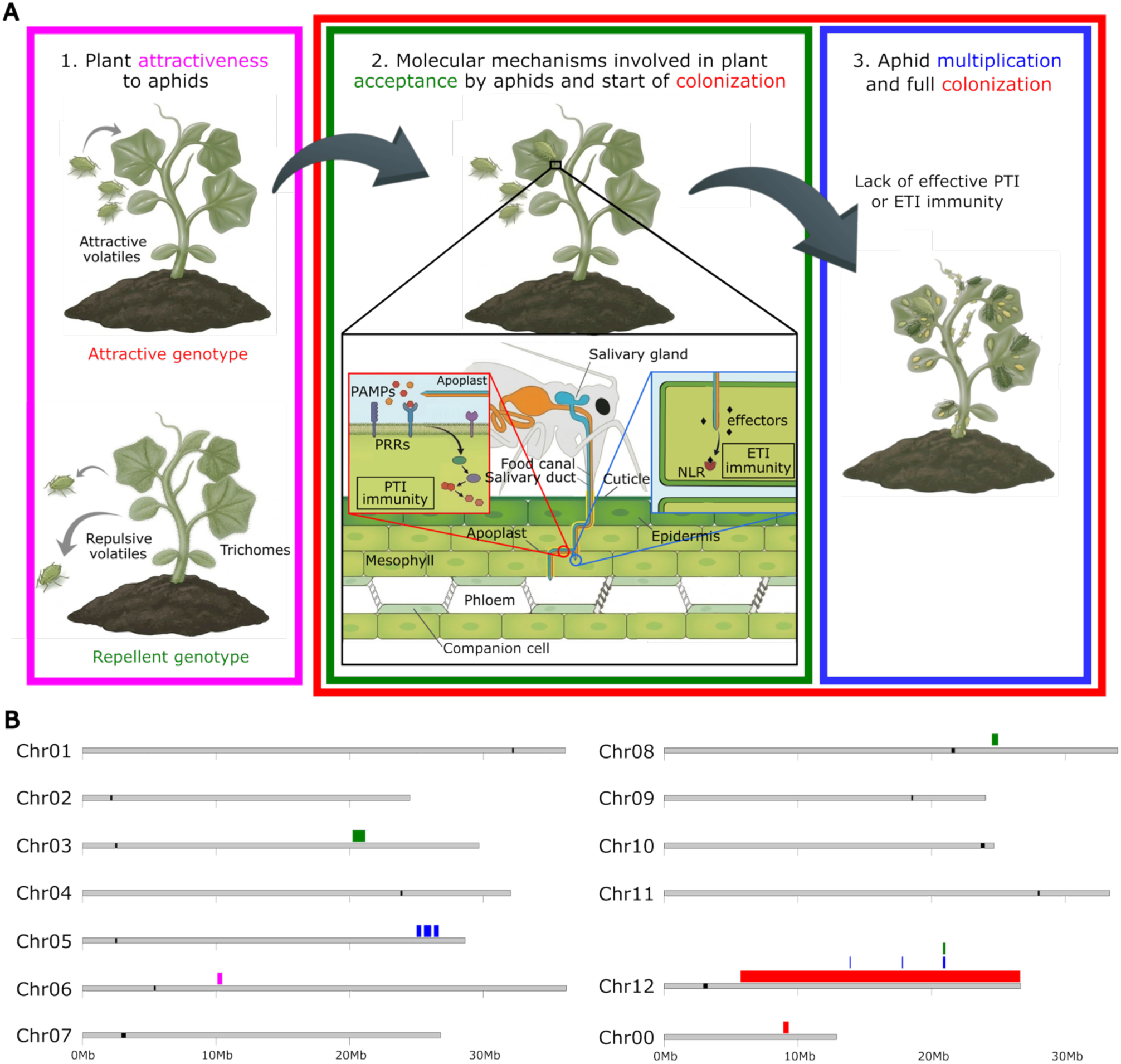
Conceptual framework of different steps of plant–aphid interactions during host colonization and their genetic control of resistance in melon. **A,** First, aphids are attracted to the plant or repelled by visual and chemical cues emitted by the host (pink section, attractiveness). After landing, aphids probe plant tissues with their stylets and initiate feeding, triggering plant defense responses that determine feeding acceptance (green section). During this phase, aphid saliva deposited in the apoplast may contain PAMP-like molecules that activate PAMP-triggered immunity (PTI), while watery saliva injected into plant cells during stylet punctures may deliver effectors that can be recognized by plant intracellular NLR receptors, leading to effector-triggered immunity (ETI). These defense layers collectively determine whether aphids can successfully colonize the plant (red section). If plant defenses are overcome, aphids establish sustained phloem feeding and reproduce, resulting in aphid multiplication (blue section). **B,** Loci controlling melon resistance to different life traits of *A. gossypii* CUC1 clone. Loci in pink, green and blue were detected by SNP-based GWAS and were linked to low attractiveness, acceptance and multiplication, respectively. Loci in red were detected by QTL-seq and were linked to low colonization.

Here, we presented a comprehensive overview of the genetic architecture underlying melon resistance to the clone of *A. gossypii* with genotype CUC1 (Fig. 8B), based on a series of complementary traits that reflect the multiple layers of the interaction between plants and aphids. Indeed, a diverse array of plant defense mechanisms influences aphid behavior, survival, development and reproduction at various stages of infestation ^[64]^. The very first layer of plant defense against aphids involves the plant’s ability to repel them, thereby influencing their behavior and host selection. Plants possess a variety of mechanisms to achieve this, including the production of volatile compounds ^[66,67]^, the development of waxy cuticles and non-glandular trichomes ^[68,69]^, or the presence of glandular trichomes that can secrete secondary metabolites capable of inducing dispersal behavior in aphids ^[70,71]^. We recorded a trait of plant attractiveness to *A. gossypii* CUC1 clone, based on the establishment of choice bioassays. We uncovered a genomic region on chromosome 6 expanding 373 kb, likely linked to aphid attractiveness (Fig. 8B). Further studies will be necessary to pinpoint the candidate genes underlying this association. The acceptance phenotype measured the capacity of plants to limit aphids’ establishment, influencing their feeding behavior, growth and development during the first stages of aphid colonization. We found QTLs associated with aphid acceptance on chromosomes 3, 8 and 12 (Fig. 8B). Among underlying candidate genes within these regions, two closely linked genes, Chr03.1000694, encoding a MYB33 transcription factor, and Chr03.1000687, encoding an ethylene insensitive 3-like 1 (EIL1) protein, stood out due to their established roles in aphid resistance across species ^[4–8]^. Aphids feed on phloem, and plants rely on phloem-based defenses (PBD) often regulated by MYB transcription factors ^[72]^. For instance, in wheat, *TaMYB19*, *TaMYB29*, and *TaMYB44* coordinate PBD against English grain aphid (*Sitobion avenae*) through regulation of β-1,3-glucan synthases and mannose-binding lectins ^[5]^, acting via crosstalk with the ethylene pathway. In chrysanthemum, *CmMYB19* and *CmMYB15* enhance lignin biosynthesis to reinforce cell walls, therefore reducing aphid colonization ^[6,7]^. In *Brassica juncea*, the MYB alleles *BjA06.GL1* and *BjB02.GL1* were recently found to control leaf trichome formation, further contributing to aphid resistance ^[4]^. In *Arabidopsis thaliana*, *AtMYB44* activates PBD against green peach aphid (*Myzus persicae*) via the regulation of ethylene insensitive 2 (EIN2), an essential gene in ethylene signaling, and both genes are required for effective responses ^[8,9]^. In our study, the close genomic linkage of MYB33 and EIL1 in melon chromosome 3 suggests a comparable defense mechanism, given that EIL1 genes have been shown to be both necessary and sufficient for the activation of ethylene-responsive genes ^[73,74]^.

A more advanced layer of plant-aphid defense involves the plant’s ability to hinder aphid survival and reproduction after their establishment. To assess this level of interaction, we evaluated colonization and multiplication phenotypes, colonization representing a composite trait that reflects both plant acceptance and aphid reproductive capacity. Using two independent and complementary methods, BSA-NGS and SNP-based GWAS, we found evidence that some resistance mechanisms limiting *A. gossypii* CUC1 multiplication are encoded on chromosome 12 (Fig. 8B). GWAS identified three QTLs on this chromosome, one of which was also linked to aphid acceptance. Complementarily, BSA-NGS linked the capacity of ANSO-77 to limit CUC1 colonization to chromosome 12, revealing a large QTL encompassing most of the genomic regions detected by GWAS. Interestingly, ANSO-77 carried the unfavorable alleles for the top SNPs within the GWAS-evidenced QTLs, suggesting that multiple loci and allelic combinations may underlie resistance to aphid multiplication on chromosome 12. Aside from chromosome 12, SNP-based GWAS on CUC1 multiplication data reported a highly supported peak overlapping the largest cluster of NLRs in melon located on chromosome 5, which contains the *Vat* region (Fig. 8B). The top-significant SNP was located 51.7 kb upstream of the M5 marker. Subsequent GWAS built on pan-genome exploration of k-mers and graph nodes across 143 accessions further refined this association, supporting the localization of resistance to CUC1 multiplication within the *Vat* region defined by markers M5 and M4.

The *Vat* locus was associated with resistance to non-persistent viruses when transmitted by aphids and mapped to linkage group V in the 1990s ^[75,76]^. Some years later, in the early 2000s, the gene *Vat* was cloned using a back-cross population derived from a cross between the susceptible accession VEDRANTAIS and the resistant accession PI 161375 ^[77]^. This gene corresponds to the homolog *Vat* 4R65aa_B in this study. *Vat* was experimentally validated using bioassays with the *A. gossypii* clone NM1, and the *Vat* region was later associated with resistance to a large number of clones ^[61,78]^. Here, we confirmed the involvement of *Vat* alleles in resistance to the CUC1 clone. Recent studies suggested a major role of the number of R65aa motifs (located in the LRR domain) in pathogen recognition ^[23,28,79]^, and phylogeny of *Vat* homologs with four R65aa motifs from 19 accessions was congruent with resistance to CMV inoculated by NM1 and C9 aphids ^[23]^. Multiple lines of evidence in our study supported that *Vat* homologs with four R65aa motifs are also key determinants of resistance to clone CUC1. Allele-based GWAS further identified *Vat* 4R65aa_B and 4R65aa_C as the most plausible candidates, and SNP-based GWAS performed after excluding accessions with candidate *Vat* 4R65aa alleles produced no signal on chromosome 5, further supporting the role of *Vat* 4R65aa homologs in resistance to *A. gossypii*.

Based on this hypothesis, we further evaluated the efficacy of *Vat* 4R65aa homologs against five clones of *A. gossypii*: C6, C9, CUC1, GWD2 and NM1. Since resistance to non-persistent viruses triggered by aphids has been demonstrated to be qualitative ^[17]^, this trait enabled us to associate 20 4R65aa *Vat* homologs with complete resistance patterns to CMV triggered by five aphid clones. This effort offers an extensive overview of allelic variation at the *Vat* locus and its relationship with phenotypic diversity, covering nearly all the diversity currently described in melon ^[27]^. *Vat*-mediated resistance to CMV is expected to be initiated by the specific detection of an aphid effector, triggering signaling pathways that elicit plant defenses against both aphids and viruses ^[17]^. In our study, no *Vat* homolog conferred resistance to CMV when transmitted by all aphid clones, confirming that the recorded phenotype was not confounded with resistance responses induced by the virus *senso stricto. Vat*-mediated resistance was shown to reduce CMV and CABYV epidemics in melon field trials conducted in southeastern France ^[19]^ while low attractiveness co-localized with CABYV delay of infection in cucumber ^[20]^. Although the effect of *Vat* alleles on persistent viruses, such as CABYV, has not been demonstrated in the laboratory, *Vat*-resistant plant genotypes may reduce CABYV epidemics by reducing *A. gossypii* colonization and limiting its access to the phloem ^[24]^. We uncovered *Vat* homologs associated with clone-specific resistance, as well as noteworthy *Vat* haplotypes such as those of PI 482398 and PI 482420, which confer resistance to CMV transmitted by five *A. gossypii* clones. Combining *Vat* homologs conferring specific resistance to aphids with sources of resistance to viruses *senso stricto* may represent an efficient approach to strengthen resistance aphid-transmitted viruses ^[80,81]^. Such combinations could enhance the durability of resistance, particularly in the event of the emergency of new vectors that do not trigger *Vat*-mediated defense. Complementarily, *Vat* homologs conferring resistance to viruses may constitute a retaining wall limiting the ability of viruses to rapidly overcome resistances to viruses *senso stricto*.

Keeping in mind the multiple layers of interaction between aphids and plants, combining resistance loci affecting different stages of this interaction could enhance the breadth and potentially the overall level of resistance. In this study, we identified QTLs associated with different components of melon resistance to the *A. gossypii* CUC1 clone, including plant attractiveness (chromosome 6), plant acceptance by aphids (chromosomes 3, 8, 12), and aphid colonization and multiplication (chromosomes 5, 12). Given the multiple GWAS performed, concordance of signals across complementary marker representations provides additional support for the robustness of the identified regions. Pyramiding these QTLs linked to different components of the interaction and located on different chromosomes could therefore provide broader and potentially stronger resistance than relying on a single mechanism. Such pyramiding may also reduce the likelihood that aphid populations can overcome resistance through adaptation to a single mechanism, although the durability of these combinations remains to be experimentally assessed. Because resistance to *A. gossypii* can be clone-specific, these findings on the genotype CUC1 should however not necessarily be extrapolated to other aphid clones. In this context, our characterization of *Vat* alleles with different specificities toward aphid clones could enable more targeted pyramiding strategies and fine breeding, taking into account the occurrence and prevalence of different aphid clones in melon production basins.

## Acknowledgements

We are grateful to CEA-IbFJ-Genoscope for providing sequencing, bioinformatics and storage facilities. We thank the Genotoul bioinformatics platform Toulouse Occitanie (Bioinfo Genotoul, https://doi.org/10.15454/1.5572369328961167E12) for providing computing and storage resources. We also thank the INRAE Pangenome Working Group PANANNOT for valuable discussions and collaborative work.

## Conflict of interest disclosure

The authors declare that they comply with the PCI rule of having no financial conflicts of interest in relation to the content of the article.

One author is recommender for one Peer Community (PCI Plants: NB).

## Data, scripts, code, and supplementary information availability

Raw ONT-NAS sequencing data were retrieved from the NCBI database under BioProject PRJNA1127998. The targeted assemblies of 21 NLR regions across 143 melon accessions, together with gene annotation and NLR annotation were retrieved from the dataset available at https://doi.org/10.57745/ZALVPU in the Recherche Data Gouv repository.

Raw short-reads previously available were retrieved from NCBI BioProjects PRJNA1164662 (for CANTON), PRJNA662717 (for ANSO-77), PRJNA662721 (for DOUBLON), PRJNA1164664 (for ZHIMALI), PRJNA1164667 (for PI 414723), PRJNA1164698 (for VEDRANTAIS), PRJNA1164660 (for ANANAS), and PRJNA1273021 (for all the rest of accessions). Short-reads data acquired during this project for the diversity panel were added to this last NCBI BioProject. Resequencing of resistant and susceptible bulks for QTL-seq were added to NCBI BioProject PRJNA1433198.

Raw phenotyping data for plant attractiveness to *A. gossypii* CUC1 clone, resistance to colonization of the ANSO-77 x VEDRANTAIS F_2_ population, resistance to colonization of the ANSO-77 x DOUBLON F_3_ population, and resistance to CMV transmitted by different clones of *A. gossypii* were deposited at the dataset available at https://doi.org/10.57745/AUO6YY in the Recherche Data Gouv repository. Raw phenotyping data for plant acceptance and aphid multiplication on the diversity panel were retrieved from CUC1 tests 239-266 from the dataset available at https://doi.org/10.57745/MDDRFF in the Recherche Data Gouv repository.

The SNP matrix for colonization data (MA_NLRome_), the pan-NLRome graph GFA file, the P/A matrix of pan-NLRome graph nodes, the k-mer P/A matrix, and the k-mer kinship matrix were retrieved from https://doi.org/10.57745/AUO6YY in the Recherche Data Gouv repository. The SNP matrix for plant attractiveness (MA_attractiveness_) was deposited at the same repository.

All scripts used for data analysis and plotting in this paper are accessible at the GitLab pages indicated in the scripts_availability.txt file included in the dataset available at https://doi.org/10.57745/AUO6YY.

## Authors’ contributions

J.B.M., A.B., and I.L. performed whole-genome short-reads sequencing. E.C. and S.M. designed and conducted multiple-choice bioassays to evaluate plant attractiveness to aphids. P.M. and K.L. managed seeds lots availability, generated the bi-parental populations, and conducted disease phenotyping of all the rest of evaluated traits. J.B.M developed and applied GWAS, pre-GWAS and post-GWAS methods, and conducted all statistical analyses. V.R.R. performed DNA extractions and genotyping of the ANSO-77 x VEDRANTAIS population. J.L. conducted bulk-segregant analysis and provided general bioinformatics expertise. J.B.M. and A.C. did the data submission. P.F.R., N.B. and D.H. conceived the study and provided expertise. J.B.M. and N.B. wrote the manuscript. All authors read and approved the final manuscript.

