## Supplementary Figures for "Linkage and association mapping coupled with pan-genome analyses of *Vat* homologs reveal QTLs and alleles for aphid resistance in melon"

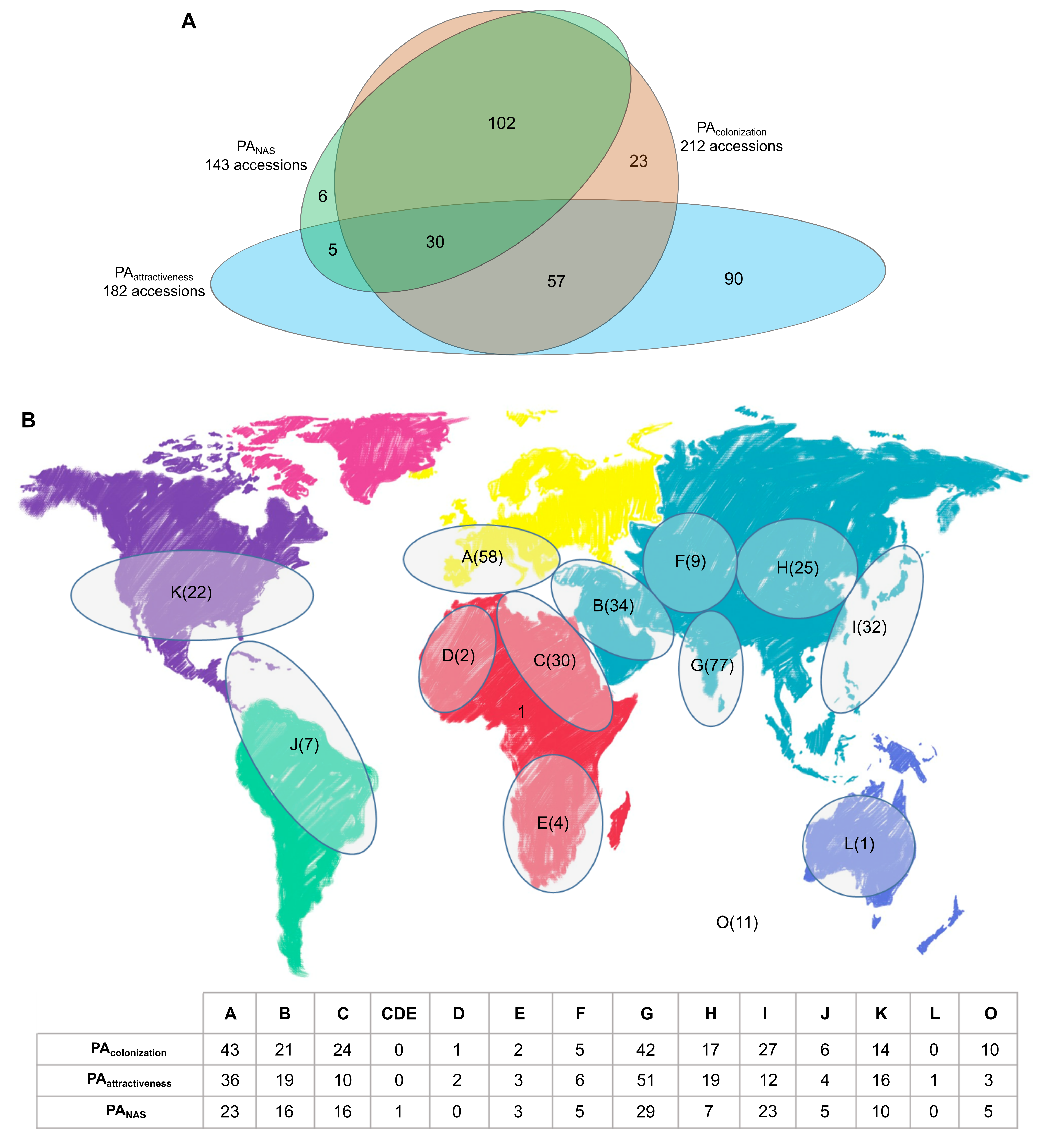

**Fig. S1.** Distribution of melon accessions across diversity panels and geographical origins. **A,** Venn diagram indicating the number of accessions composing the PA_colonization_, PA_attractiveness_ and PA_NAS_ diversity panels, as well as the number of accessions shared among panels. **B,** Geographical distribution of the accessions used in this study. The total number of accessions per geographical region is depicted on the world map. The table below details the distribution of accessions among the different diversity panels within each region. Each geographical origin is represented by a distinct letter; O indicates accessions with unknown origin.

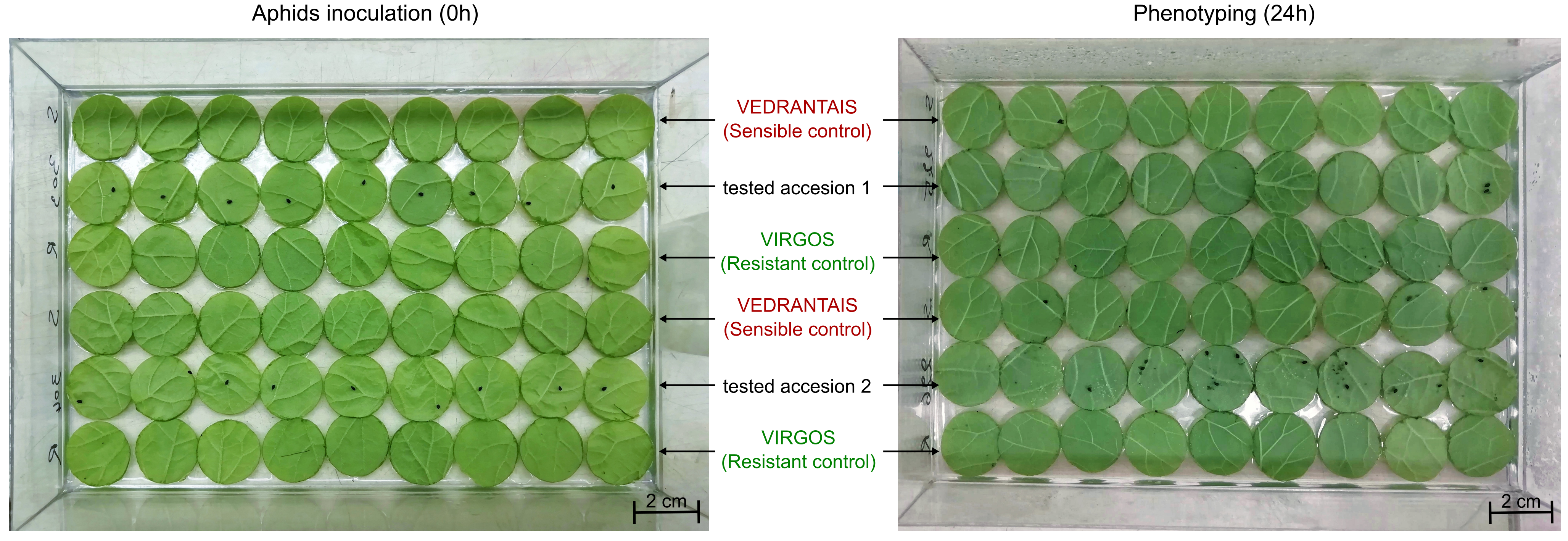

**Fig. S2**. Experimental layout of genotype-specific leaf disks and aphid placement for attractiveness bioassays. Representative images taken at the time of aphid placement (0 h) and at phenotyping (24 h post-inoculation) are shown on the left and right sides, respectively. The two images do not correspond to the same experimental box.

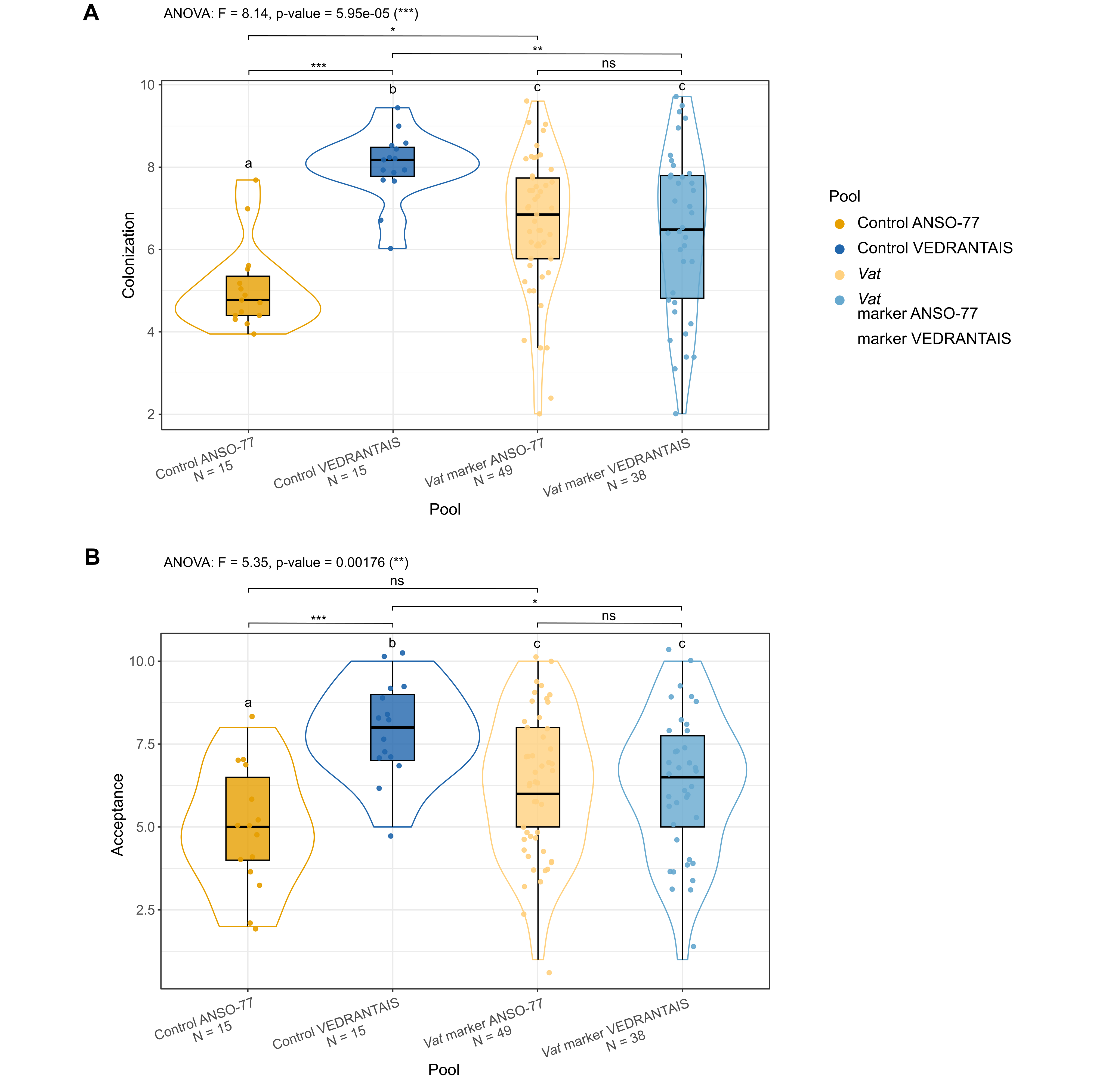

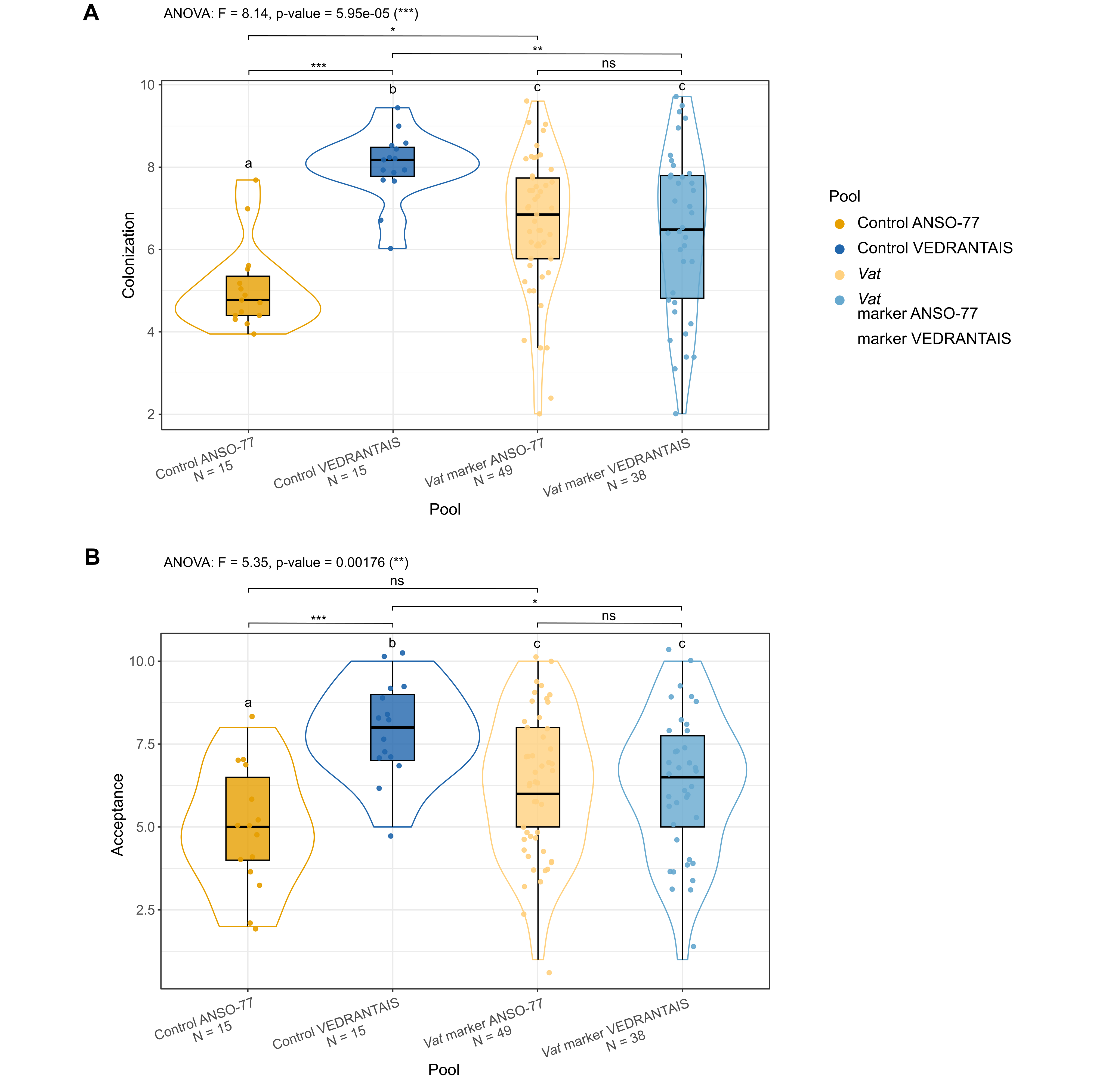

**Fig. S3.** Distribution of *A. gossypii* CUC1 colonization **(A)** and acceptance **(B)** in selected F_2_ pools from the ANSO-77 x VEDRANTAIS cross. We isolated genomic DNA from 186 F_2_ plantlets and genotyped by PCR using the markers Z5259F and Z5238R (specific of the *Vat* cluster of ANSO-77) and Z1431F and Z5239R (specific of the *Vat* cluster of VEDRANTAIS). Based on amplification profiles, 38 F_2_ plants displaying the VEDRANTAIS pattern and 49 displaying the ANSO-77 pattern were selected for phenotyping of *A. gossypii* CUC1 acceptance and multiplication. ANSO-77 and VEDRANTAIS controls (15 plants) are also shown. Differences among groups were assessed by one-way ANOVA followed by Tukey’s HSD post-hoc test. Tukey’s HSD groups are marked with letters, and pairwise significance levels are denoted as ns (p ≥ 0.05), * (p < 0.05), ** (p < 0.01), and *** (p < 0.001).

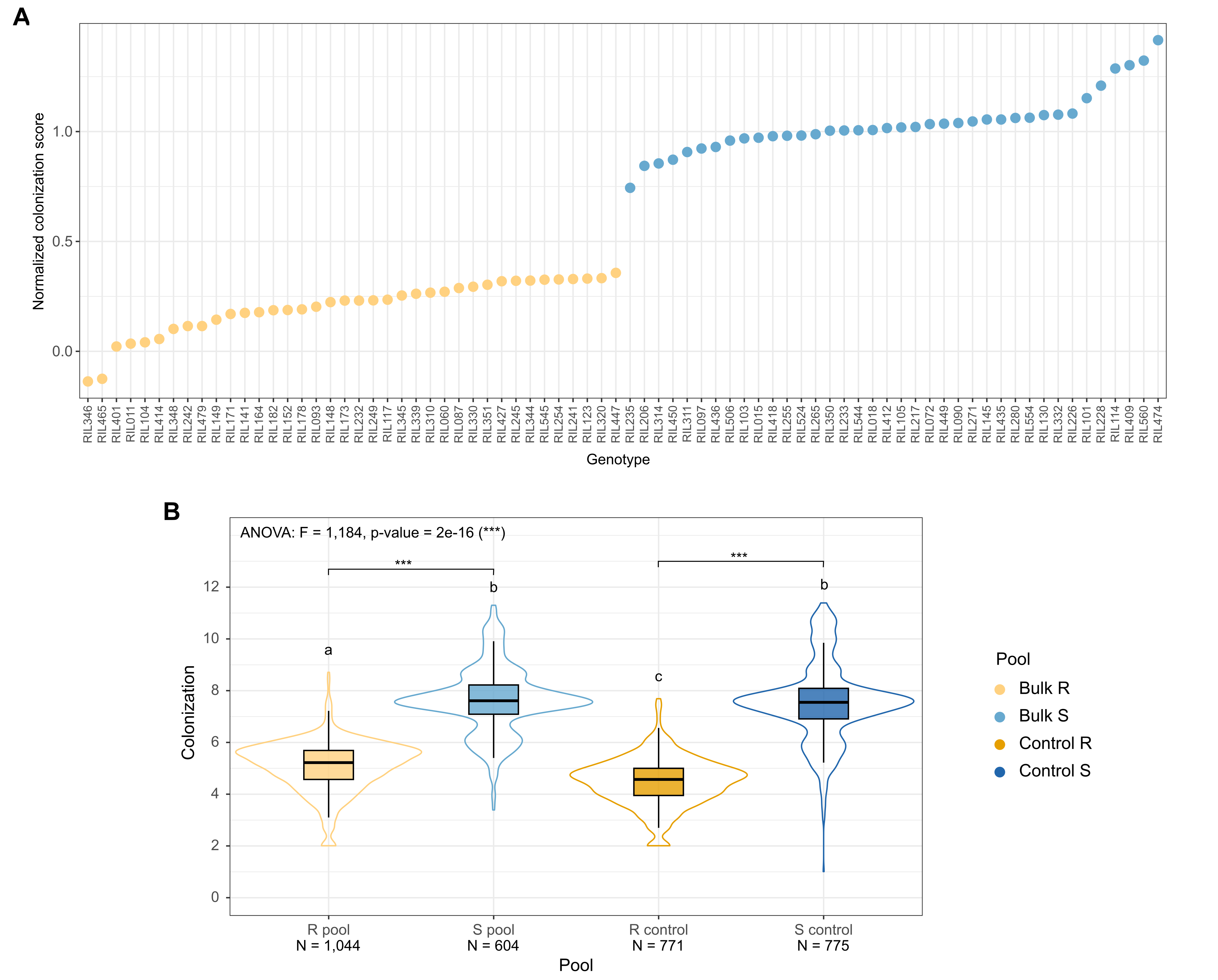

**Fig. S4.** Colonization of *A. gossypii* CUC1 clone on 76 selected F_3_ families derived from ANSO-77 x DOUBLON cross. **A,** Mean and standard deviation of *A. gossypii* CUC1 colonization for each of the 76 selected F_3_ families. Mean values of families included in the resistant (R) and susceptible (S) bulks are indicated as light orange and light blue points, respectively. Resistant (VIRGOS) and susceptible (VEDRANTAIS) controls are depicted with dark orange and dark blue points, respectively. We tested at least 15 individuals for most families. **B,** Distribution of *A. gossypii* CUC1 colonization in resistant and susceptible F_3_ pools and in the parental controls. The number of plants phenotyped on each group (N) is depicted at the bottom of the plot. Differences among groups were tested using one-way ANOVA followed by Tukey’s HSD post-hoc test. Tukey’s HSD groups are marked with letters, and pairwise significance levels between the two controls and between the two pools are denoted as ns (p ≥ 0.05), * (p < 0.05), ** (p < 0.01), and *** (p < 0.001).

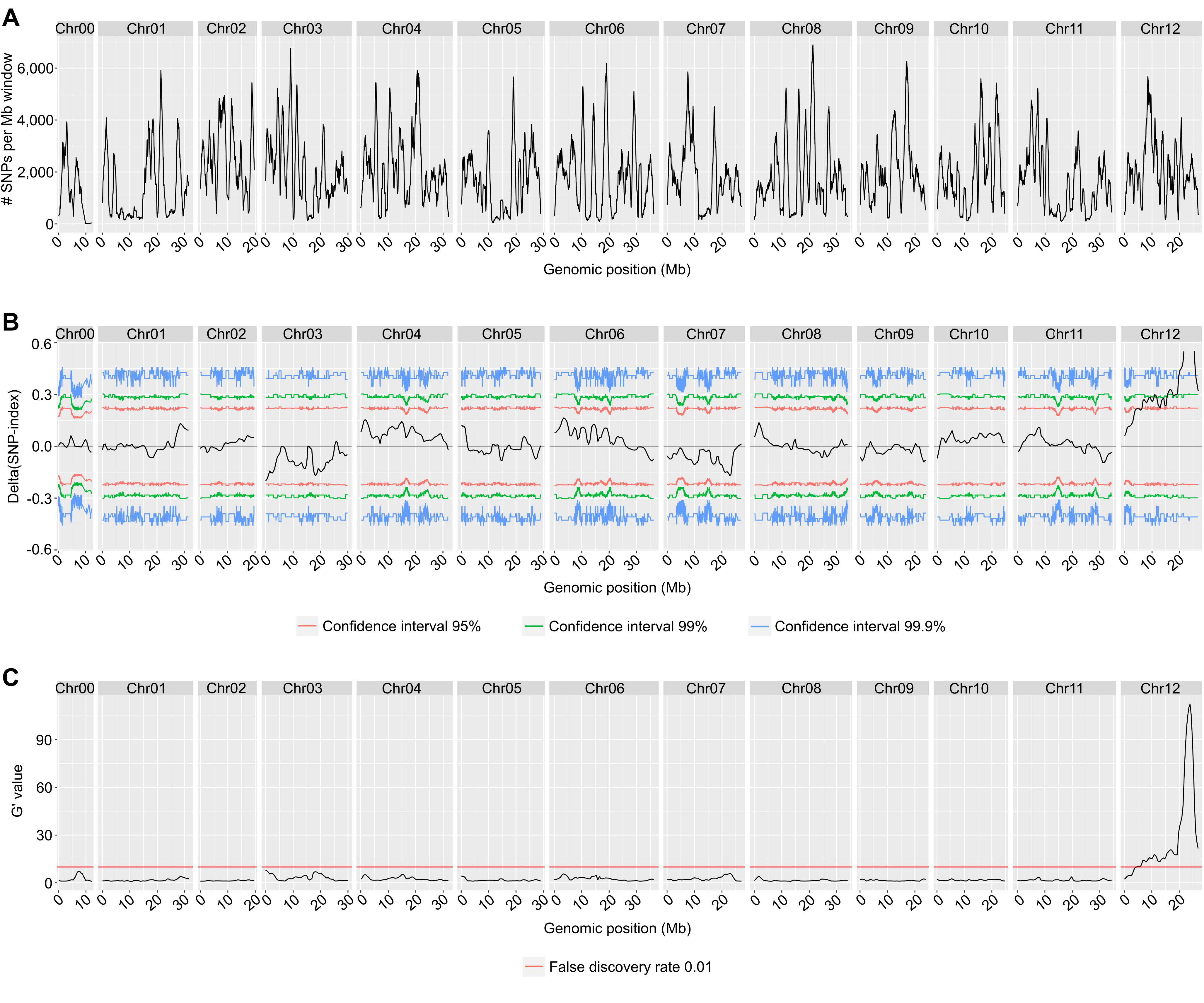

**Fig. S5.** QTLs associated with resistance to *A. gossypii* clone CUC1 colonization, identified by QTL-seq using the DOUBLON genome as reference. All analyses were performed using a 1 Mb sliding window. **A,** Distribution of SNPs and short INDELs within each 1 Mb smoothing window along the 12 DOUBLON chromosomes and unassembled contigs. **B,** Tricube-smoothed Δ(SNP-index) distribution along the 12 DOUBLON chromosomes and unassembled contigs. Red, green and blue lines indicate the 95%, 99% and 99.9% two-sided confidence intervals, respectively. **C,** Tricube-smoothed G’ values distribution along the 12 DOUBLON chromosomes and unassembled contigs. The red horizontal line represents a genome-wide false discovery rate threshold of 0.01.

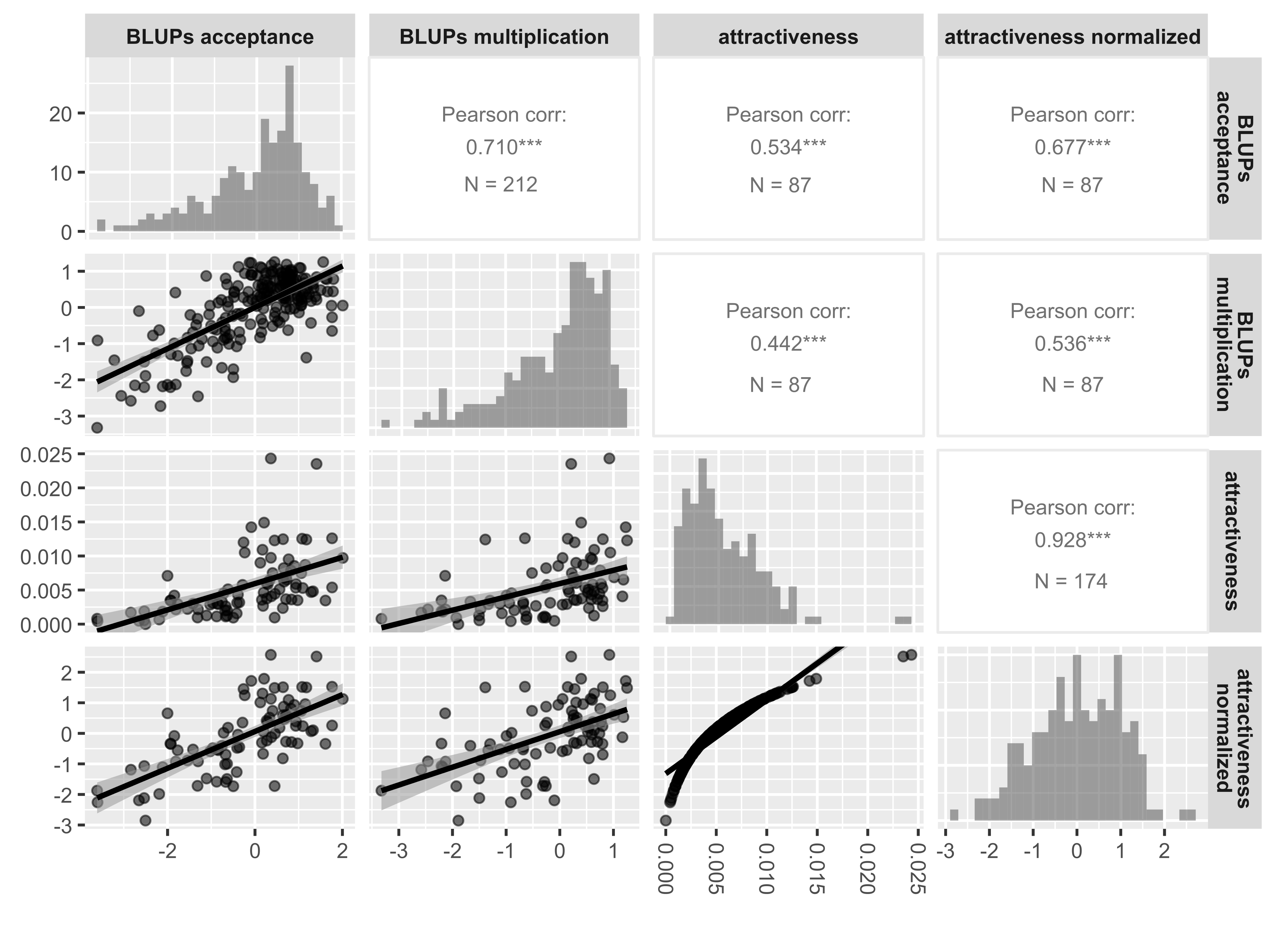

**Fig. S6.** Correlation matrix of phenotypic traits related to plant response to *A. gossypii* CUC1. Pairwise relationships are displayed as scatterplots in the lower triangle, trait distributions are displayed along the diagonal, and Pearson correlation coefficients are presented in the upper triangle. Significance is coded as * (p < 0.05), ** (p < 0.01), and *** (p < 0.001). N values indicate the number of shared accessions used for each pairwise comparison.

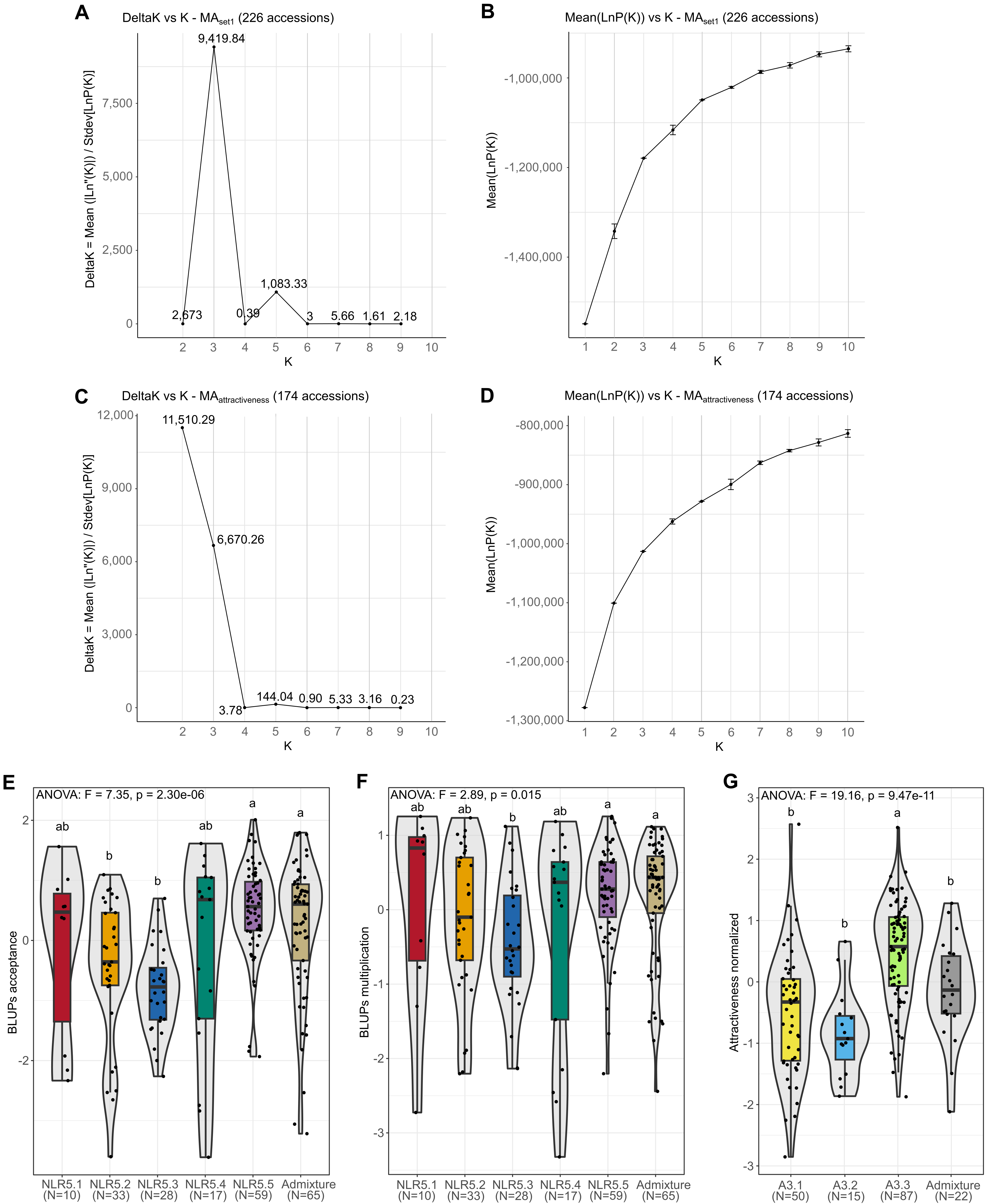

**Fig. S7.** Population structure inferred by STRUCTURE analysis and distribution of phenotypes across inferred genetic groups. **A and C,** Evanno plots showing Delta K values for ten tested K values (population structured into K genetic groups), averaged across ten independent replicates per K. Delta K distribution suggested that the set1 population (226 accessions, including the 212 composing the PA_colonization_ panel) is most likely structured into three or five genetic groups (A), whereas the attractiveness population (174 accessions phenotyped for plant attractiveness to *A. gossypii* CUC1 clone) is most likely structured into two or three genetic groups (C). **B and D,** Distribution of mean LnP(K) values (log probability of the data) ± standard deviation across ten replicates per K value. Similar to (A) and (C), LnP(K) supported a most probable structure of three or five genetic groups for the set1 population, and two or three genetic groups for the attractiveness population. **E, F and G,** Distribution of BLUPs for aphid acceptance (E), aphid multiplication (F), and normalized plant attractiveness across five (E and F) and three (G) inferred genetic groups. Accessions showing admixture between groups are plotted as a separate category. Dots represent individual accessions. The number of phenotyped accessions (N) per genetic group is indicated at the bottom of each plot. Differences among groups were tested using one-way ANOVA followed by Tukey’s HSD post-hoc test.

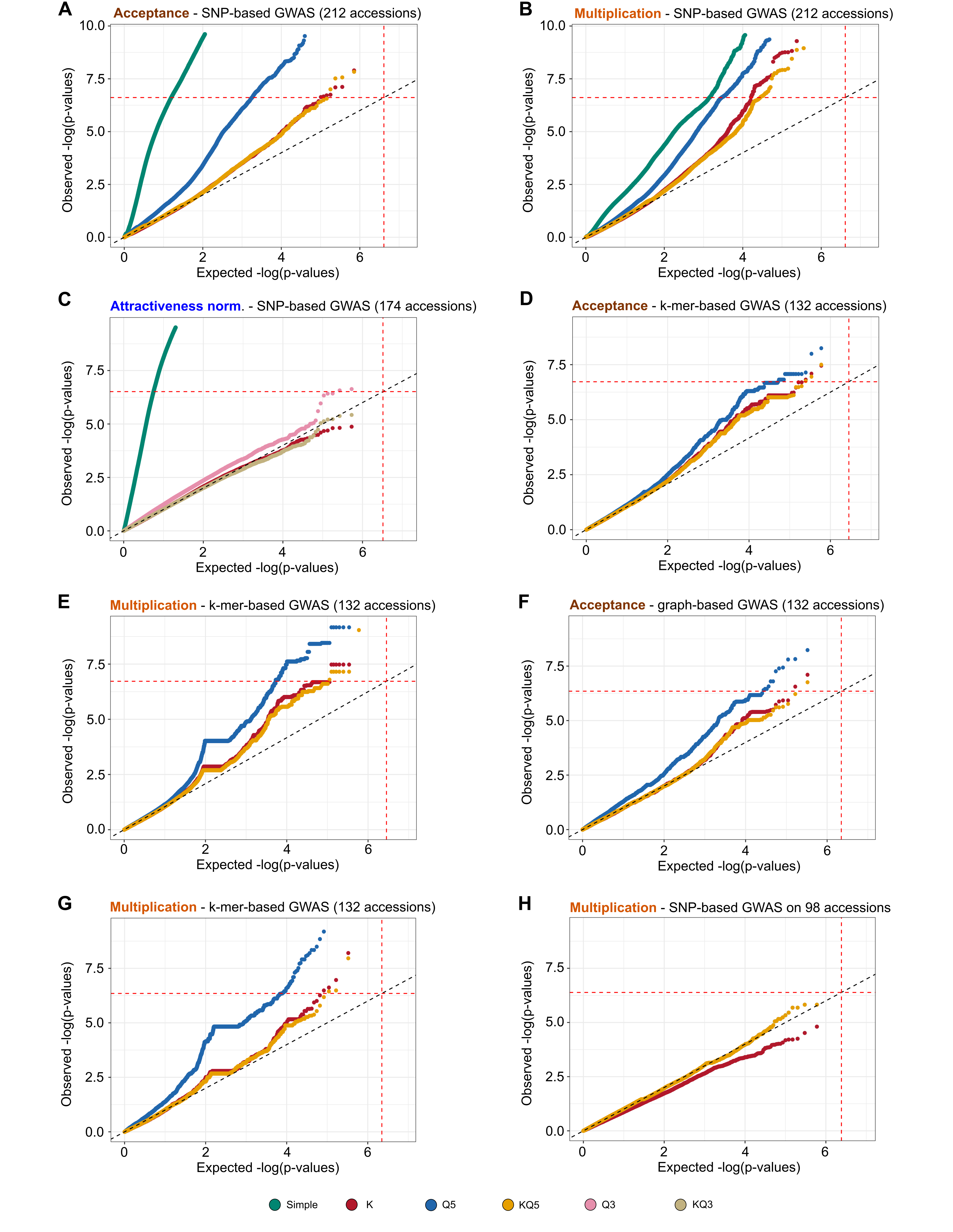

**Fig. S8.** Quantile-quantile (Q-Q) plots for selection of the best-fit GWAS model. **A and B**, Q-Q plots from SNP-based GWAS for *A. gossypii* CUC1 clone acceptance (A) and multiplication (B). **C,** Q-Q plot from SNP-based GWAS for plant attractiveness to the *A. gossypii* CUC1 clone. **D and E,** Q-Q plots from k-mer-based GWAS for response to CUC1 acceptance (D) and multiplication (E). **F and G,** Q-Q plots from graph-based GWAS for response to CUC1 acceptance (F) and multiplication (G). **H,** Q-Q plot from SNP-based GWAS for multiplication using a reduced panel of 98 accessions for which the NLRome was assembled and that lack *Vat* homologs carrying four R65aa motifs likely associated with CUC1 resistance. In all analyses, we compared four GWAS models: a simple model without correction for relatedness or population structure, a kinship-only model (K), a model accounting for population structure (Q), and a combined model including both kinship and structure (KQ). Q-Q plots were generated under the assumptions of the random regression model. Colored points show the -log10(p-values) of tested SNPs under different models. The diagonal dashed lines indicate the expected -log10(p-values) under the null hypothesis of no association. The red dashed lines indicate the Bonferroni-corrected significance thresholds, calculated as α = 0.05 divided by the number of estimated independent SNPs.

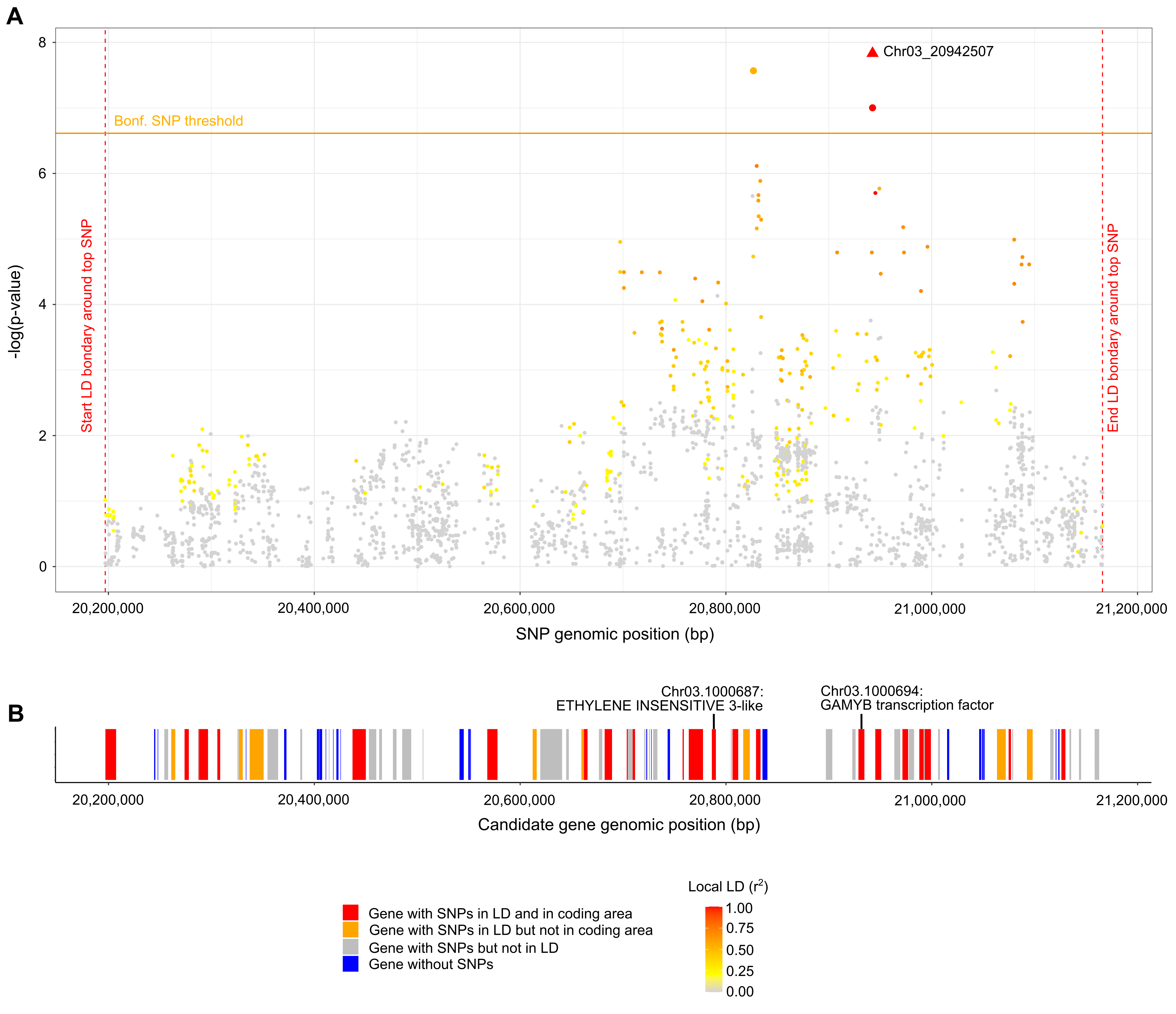

**Fig. S9.** Insights into the QTL on chromosome 3 detected through SNP-based GWAS for *A. gossypii* CUC1 acceptance. **A,** Manhattan plot of SNPs located within the QTL interval on chromosome 3. The QTL boundaries (red vertical dotted lines) were defined based on LD decay around the top-significant SNP (Chr03_20942507; red triangle). A 100 kb window centered on this SNP was iteratively expanded until no SNPs remained with an r² > 0.2 relative to Chr03_20942507. Each dot represents an SNP which is colored according the local LD relative to Chr03_20942507. The horizontal orange line indicates the Bonferroni-corrected significance threshold, calculated with α = 0.05, adjusted for the estimated number of independent variants. **B,** Genes located within the represented QTL, with top candidate genes in red (harboring at least one SNP in LD with the top SNP and located in a coding area), potential candidate genes in orange (harboring at least one SNP in LD with the top SNP and located in a non-coding area), and other genes in gray (harboring SNPs but not in LD with the top SNP) or blue (not harboring SNPs).

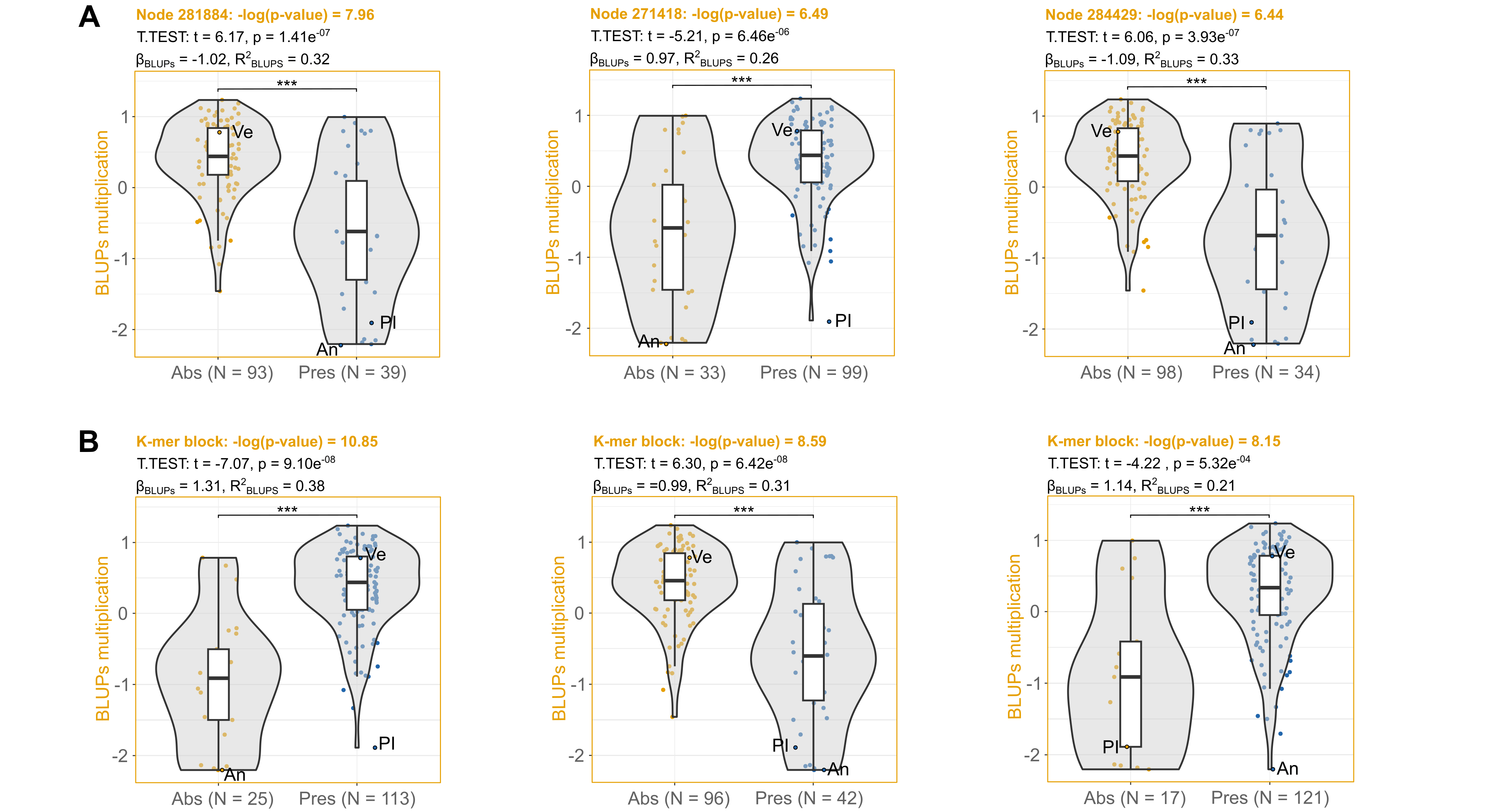

**Fig. S10.** Allelic effects of pan-NLRome graph nodes **(A)** and k-mers **(B)** associated with plant responses to *A. gossypii* clone CUC1 multiplication. Box-plots and violin plots represent the distribution of BLUPs across accessions carrying (pres.) or lacking (abs.) each reported variant. Dots indicate phenotypic values of individual accessions. Ve: VEDRANTAIS (susceptible); An: ANSO-77 (resistant); PI: PI 414723 (resistant). The number of accessions contributing to each group is indicated at the bottom of the graphs (N). Differences between groups were tested using a t-test, with significance levels denoted as * (p < 0.05), ** (p < 0.01), and *** (p < 0.001). Linear regression coefficients (β_BLUPs_) indicate the effect size of the presence of each variant, and R² values indicate the proportion of phenotypic variance explained.

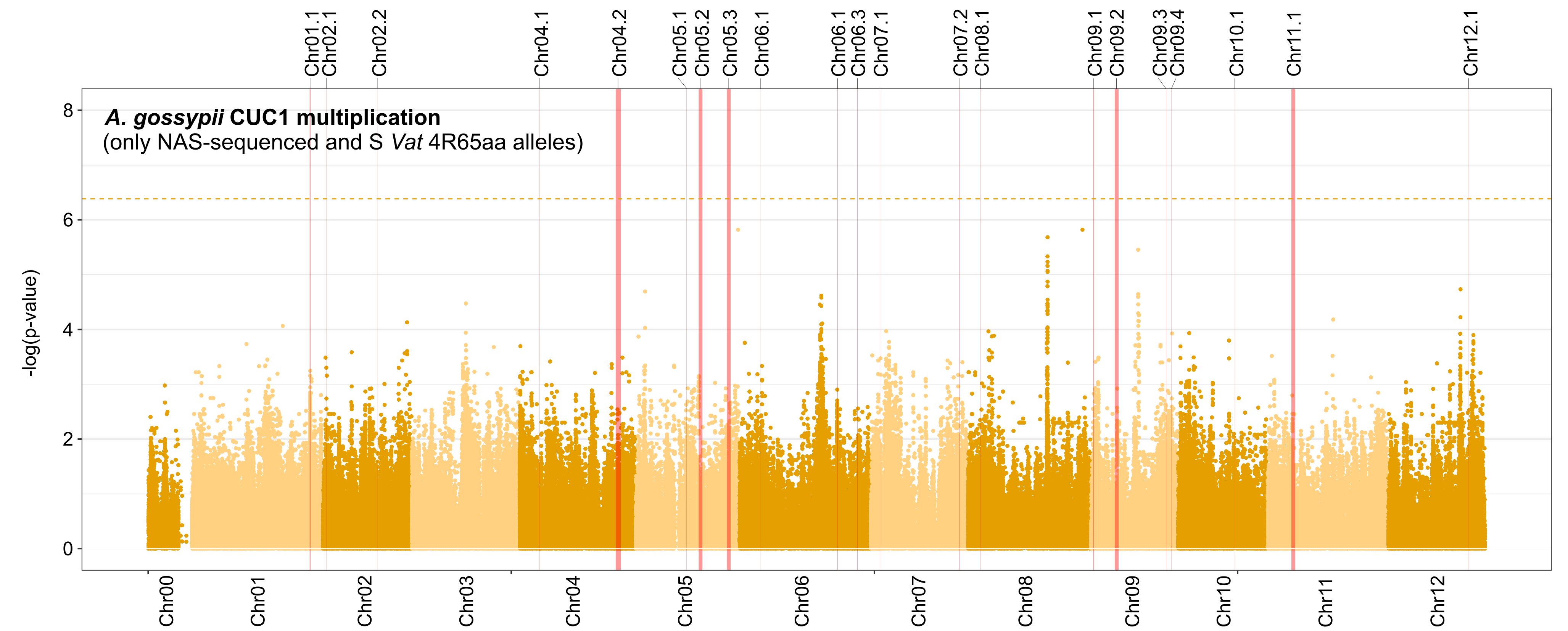

**Fig. S11.** GWAS results for *A. gosypii* CUC1 multiplication in a reduced panel of 98 accessions. These accessions have their NLRome assembled and lack *Vat* homologs with four R65aa motifs likely associated with CUC1 resistance. P-values were obtained from the first MLMM step. The horizontal dashed line indicates the Bonferroni-corrected significance threshold, calculated with α = 0.05, adjusted for the estimated number of independent variants. Vertical red bars locate NLR-containing genomic regions, with bar length scaled to their sizes.
